# Unusual mode of dimerization of retinitis pigmentosa-associated F220C rhodopsin

**DOI:** 10.1101/2020.12.28.424580

**Authors:** George Khelashvili, Anoop Narayana Pillai, Joon Lee, Kalpana Pandey, Alexander M. Payne, Zarek Siegel, Michel A. Cuendet, Tylor R. Lewis, Vadim Y. Arshavsky, Johannes Broichhagen, Joshua Levitz, Anant K. Menon

**Affiliations:** Department of Physiology and Biophysics, Weill Cornell Medical College, New York, NY, 10065; Institute of Computational Biomedicine, Weill Cornell Medical College, New York, NY 10065; Department of Biochemistry, Weill Cornell Medical College, New York, NY, 10065; Tri-Institutional PhD Program in Chemical Biology, Weill Cornell Medical College, New York, NY, 10065; Neurosciences Graduate Program, University of California San Diego, La Jolla, CA 92093; Ludwig Institute for Cancer Research, University of Lausanne, and Department of Oncology, University Hospital of Lausanne, 1009, Lausanne, Switzerland; Swiss Institute of Bioinformatics, 1015 Lausanne, Switzerland; Department of Ophthalmology, Duke University Medical Center, Durham, NC, 27710; Leibniz-Forschungsinstitut für Molekulare Pharmakologie, Department of Chemical Biology, Robert-Rössle-Str. 10, 13125 Berlin, Germany

**Keywords:** dimerization, fluorescence resonance energy transfer (FRET), G protein-coupled receptor (GPCR), membrane protein, molecular dynamics, phospholipid scramblase, retinitis pigmentosa

## Abstract

Mutations in the G protein-coupled receptor (GPCR) rhodopsin are a common cause of autosomal dominant retinitis pigmentosa, a blinding disease. Rhodopsin self-associates in the membrane, and the purified monomeric apo-protein opsin dimerizes *in vitro* as it transitions from detergent micelles to reconstitute into a lipid bilayer. We previously reported that the retinitis pigmentosa-linked F220C opsin mutant fails to dimerize *in vitro*, reconstituting as a monomer. Using fluorescence-based assays and molecular dynamics simulations we now report that whereas wildtype and F220C opsin display distinct dimerization propensities *in vitro* as previously shown, they both dimerize in the plasma membrane of HEK293 cells. Unexpectedly, molecular dynamics simulations show that F220C opsin forms an energetically favored dimer in the membrane when compared with the wild-type protein. The conformation of the F220C dimer is unique, with transmembrane helices 5 and 6 splayed apart, promoting widening of the intracellular vestibule of each protomer and influx of water into the protein interior. FRET experiments with SNAP-tagged wild-type and F220C opsin expressed in HEK293 cells are consistent with this conformational difference. We speculate that the unusual mode of dimerization of F220C opsin in the membrane may have physiological consequences.

## Introduction

The visual pigment rhodopsin is both a G protein-coupled receptor (GPCR) and a critical structural component of the outer segment of photoreceptor cells (1). Its unexpected phospholipid scramblase activity, evinced on reconstitution of purified protein into lipid vesicles, accounts for the lipid scrambling observed in isolated photoreceptor disc membranes (2–4). Rhodopsin can activate the visual signaling cascade as a monomer, but, like other Class A GPCRs (5,6), it self-associates to form dimers and higher order multimers. The quaternary structure of rhodopsin has been proposed to be important for its trafficking to the outer segment, structural contributions to disc formation, and phototransduction properties (6). Rhodopsin dimers have been visualized *in situ* using atomic force microscopy of photoreceptor disc membranes adsorbed on mica (7), and cryo-electron tomography using retinas cryofixed by high-pressure freezing (8). Consistent with these results, chemical crosslinking studies of native disc membranes revealed a symmetric dimer interface mediated by transmembrane helix 1 (TM1) contacts (9), and site-directed cysteine mutagenesis studies implicated transmembrane helices 4 and 5 (TM4 and TM5) in mediating dimerization of the apo-protein opsin expressed in COS1 cells (10). The involvement of these TM helices in mediating dimer formation is supported by a cryo-electron microscopy study of rhodopsin reconstituted into nanodiscs (11), as well as a report showing that synthetic TM peptides can disrupt rhodopsin dimers in both discs and HEK cells (12).

Mutations in rhodopsin are the most common cause of autosomal dominant retinitis pigmentosa (RP), a blinding disease that afflicts more than 1.5 million individuals world-wide (13–15). We discovered that F45L, V209M and F220C rhodopsin mutants that were identified in patients with RP appear to function normally as visual pigments and phospholipid scramblases but, unlike wild type rhodopsin, they fail to dimerize *in vitro* during detergent removal *en route* to reconstitution into lipid vesicles (16). We proposed that their inability to dimerize might explain their association with retinal disease. Surprisingly, biophysical studies of the V209M and F220C mutants expressed in COS cells indicate that they self-associate similarly to wild-type rhodopsin while exhibiting a minor trafficking defect (17). Furthermore, knockin mice bearing the F45L mutation are aphenotypic, while those with the F220C mutation display a small abnormality in outer segment morphology (18). Given these disparate data, we decided to analyze the F220C mutant in more detail using a multi-pronged approach to characterize the protein. We used *in vitro* and cell-based fluorescence assays to verify and extend previous data, and computational approaches based on molecular dynamics (MD) simulations of the protein in detergent micelles and in the membrane. In all our experiments we compared the behavior of F220C opsin with that of the wild-type (WT) protein. As the F220C mutation is located on TM5, we also considered more generally the role of this helix in rhodopsin dimerization.

We now report that both WT and F220C opsin dimerize in the plasma membrane of HEK293 cells, but are monomeric when purified after solubilization of the cells in the detergent dodecyl-β-maltoside (DDM). The latter observation is consistent with our MD simulations showing that opsin dimerization is energetically unfavored in a DDM-micellar environment. We developed a protocol to mimic early steps of *in vitro* membrane reconstitution by treating DDM-solubilized opsins with detergent-adsorbing resin in the absence of phospholipids. With this protocol we find that DDM-solubilized WT opsin monomers self-associate on partial removal of detergent, whereas F220C opsin remains monomeric. Thus, consistent with previous results (16,17), the two proteins exhibit different behaviors in a micellar environment, whereas both form dimers when they are located in a membrane bilayer. These findings support the idea that WT and F220C opsins have distinct dimerization properties, motivating a deeper analysis in the membrane context.

MD simulations showed that F220C opsin forms an energetically favored dimer in the membrane when compared with the WT protein. The conformation of the F220C dimer is unique, with the TM5/TM6 helices tilted to form a splayed, criss-crossed structure. The difference in conformation of WT and F220C dimers in the membrane is supported by experiments showing that the two homo-dimers exhibit different extents of Förster resonance energy transfer (FRET) in the membrane. The splayed conformation of F220C opsin dimers promotes widening of the intracellular vestibule of each monomer in the dimer enabling water to enter the protein interior. Further analyses reveal that this unique dimerization pose is a feature of the F220C mutation and is not observed in simulations in which other positions along the TM5 interface are mutated. We speculate that the unusual mode of dimerization of F220C opsin in the membrane may underscore its physiological consequences.

## Results and Discussion

### Dimerization of WT opsin, but not F220C opsin, in a micellar system

We previously proposed that whereas both WT and F220C opsins are monomers when purified from DDM-solubilized cells, gradual withdrawal of detergent causes WT opsin but not F220C opsin to dimerize (3,16). This proposal was supported by experiments showing that immobilized WT opsin could pull-down soluble WT opsin as DDM was reduced from 1.96 to 0.2 mM, just above its critical micellar concentration (cmc)(0.15 mM), whereas the same did not hold for the F220C construct (16). Here we deploy fluorescence measurements to test key aspects of this proposal.

To establish that WT and F220C opsins are monomers when purified, we used Single Molecule Pulldown (SiMPull) (19–21). We expressed HA-SNAP-tagged WT and F220C opsins in HEK293T cells, and labeled the cell surface population with the membrane-impermeant SNAP dye LD555 before solubilizing the cells in 1% (w/v) DDM to extract the labeled proteins. The samples were subsequently transitioned to a lower detergent concentration (0.1% (w/v) DDM) and immobilized on a polyethylene glycol (PEG)-passivated coverslip via anti-HA antibodies. Individual fluorescent spots were visualized using total internal reflection (TIRF) microscopy (Figure 1A, C). For both opsin constructs, most spots were bleached within 30 s of the onset of laser illumination (representative traces shown in Figure 1A, C), and bleaching analysis revealed that ~90% of spots bleached in a single step (Figure 1B, D). These results indicate that both WT and F220C opsins are monomers in 0.1% (w/v) DDM.

**Figure 1.**
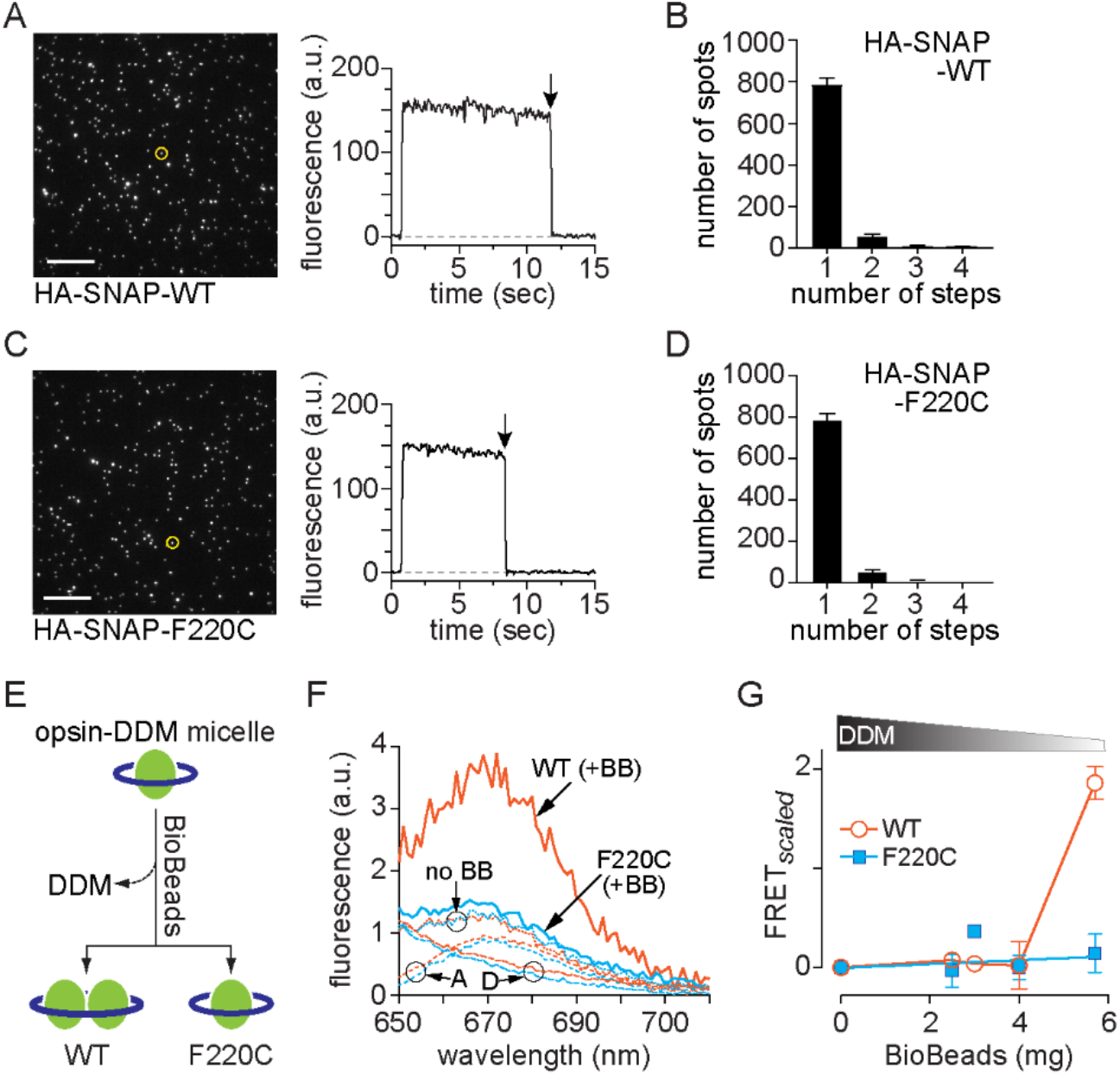
Dimerization of WT opsin, but not F220C opsin, in a micellar system. **(A)-(D)** SiMPull experiments with HA-SNAP-WT opsin (A-B) and HA-SNAP-F220C opsin (C-D). Opsin constructs were expressed in HEK293T cells and labeled using a membrane-impermeant SNAP fluorophore (LD555). The cells were solubilized in DDM and the proteins were captured on glass slides coated with anti-HA antibodies for visualization of fluorescence by TIRF microscopy (A and C). Bleaching of individual spots was recorded; traces for representative spots (circled in the fluorescence images) are shown, with arrows indicate bleaching step) and the number of spots that bleached in 1, 2, 3 or 4 steps was quantified (B and D). **(E)** Schematic of the effect of detergent removal on the quaternary structure of WT and F220C opsins. **(F)** Emission spectra (λ_ex_ =555 nm) for donor (D)-acceptor (A) pairs of WT (orange) or F220C opsin (blue) treated with 5.7 mg BioBeads (+BB, solid lines) or mock-treated (no BB, dotted lines). Spectra of mock treated samples of acceptor alone and donor alone (‘A’ and ‘D’, dashed lines) are also shown. Sensitized emission of the acceptor (λ_ex_ =555 nm, λ_em_ =670 nm) indicates FRET as seen in the BioBead-treated WT sample (+BB). **(G)** FRET was quantified after extracting increasing amounts of DDM (shown schematically above the graph) by treating samples with different amounts of BioBeads. A FRET measure (FRET_*scaled*_) was obtained by calculating the ratio of the emission intensity at 670 nm for D-A mixtures relative to the acceptor-only sample, followed by an offset correction corresponding to mock-treated samples (no BB, panel A). Data are mean ± SEM (n≥3) and connecting lines are drawn to guide the eye.

We next used Förster resonance energy transfer (FRET) to test the ability of the proteins to self-associate as a function of detergent removal in the absence of lipid vesicles (Figure 1E). For this, we fluorescently labeled purified SNAP-tagged opsins with either SNAP-Surface 549 (donor (D)) or SNAP-Surface 649 (acceptor (A)) fluorophores. Equimolar mixtures of D and A-labeled WT opsin (in 0.1% (w/v) DDM (1.96 mM)) were mock-treated, or treated with different amounts of BioBeads (BB) to reduce detergent levels. F220C opsin samples were analyzed in parallel. Fluorescence emission spectra (Figure 1F) indicate FRET in the WT opsin mixture treated with 5.7 mg BioBeads (WT+BB), but not in the F220C opsin sample (F220C+BB). No FRET was seen in mock-treated samples (dotted lines, no BB (Figure 1F)). Based on our previous calibration of BB-mediated detergent removal (3), the amount of DDM remaining after treatment of the sample with 4-6 mg BB remains above cmc. Quantification of FRET as a function of BioBead treatment, i.e. amount of detergent removed, shows that WT opsin, but not F220C opsin, dimerizes/multimerizes in samples treated with >4 mg BioBeads (Figure 1G). These results support the mechanism presented above and shown schematically in Figure 1E, i.e. that the F220C mutation prevents opsin from dimerizing as detergent is withdrawn.

### Computational analysis of the behavior of WT and F220C opsin in DDM micelles

Hypothesizing that the different behavior of WT and F220C opsin in detergent would be reflected in differences in the organization of the detergent micelle surrounding these proteins, we carried out MD simulations of opsin monomers and dimers in DDM micelles. We used a detergent: protein ratio of 150:1, based on a previous estimate of the size of DDM micelles (22,23). As shown in Figure 2 (top two rows), the organization of DDM molecules around monomeric WT and F220C opsins is as expected, with hydrophobic tails directed towards the protein, shielding the hydrophobic core of the protein from unfavorable exposure to the solvent. However, the arrangement of detergent around the dimeric constructs is strikingly different. Thus, as illustrated in Figure 2 (bottom two rows), the dimer interface of both WT and F220C systems is exposed to DDM headgroups, as the detergent molecules align vertically, akin to lipids. Such an arrangement results in the exposure of polar DDM headgroups to hydrophobic residues at the dimer interface leading to an energetically costly hydrophobic mismatch between the protein dimer and the micelle. We calculated the residual hydrophobic mismatch (RHM) energy for each protein sidechain. The concept of RHM energy and its relation to protein-protein interactions in lipid membranes has been well established (see Refs. (24,25)). It has been shown that the cost of RHM can be mitigated by modes of protein association in the bilayer that would cover the sites of RHM and eliminate the costly interaction of the protein with the membrane (26,27). Here, following the same concept, we analyzed RHM between the protein residues and DDM molecules in order to quantify the contribution of hydrophobic mismatch to the dimerization energy.

**Figure 2.**
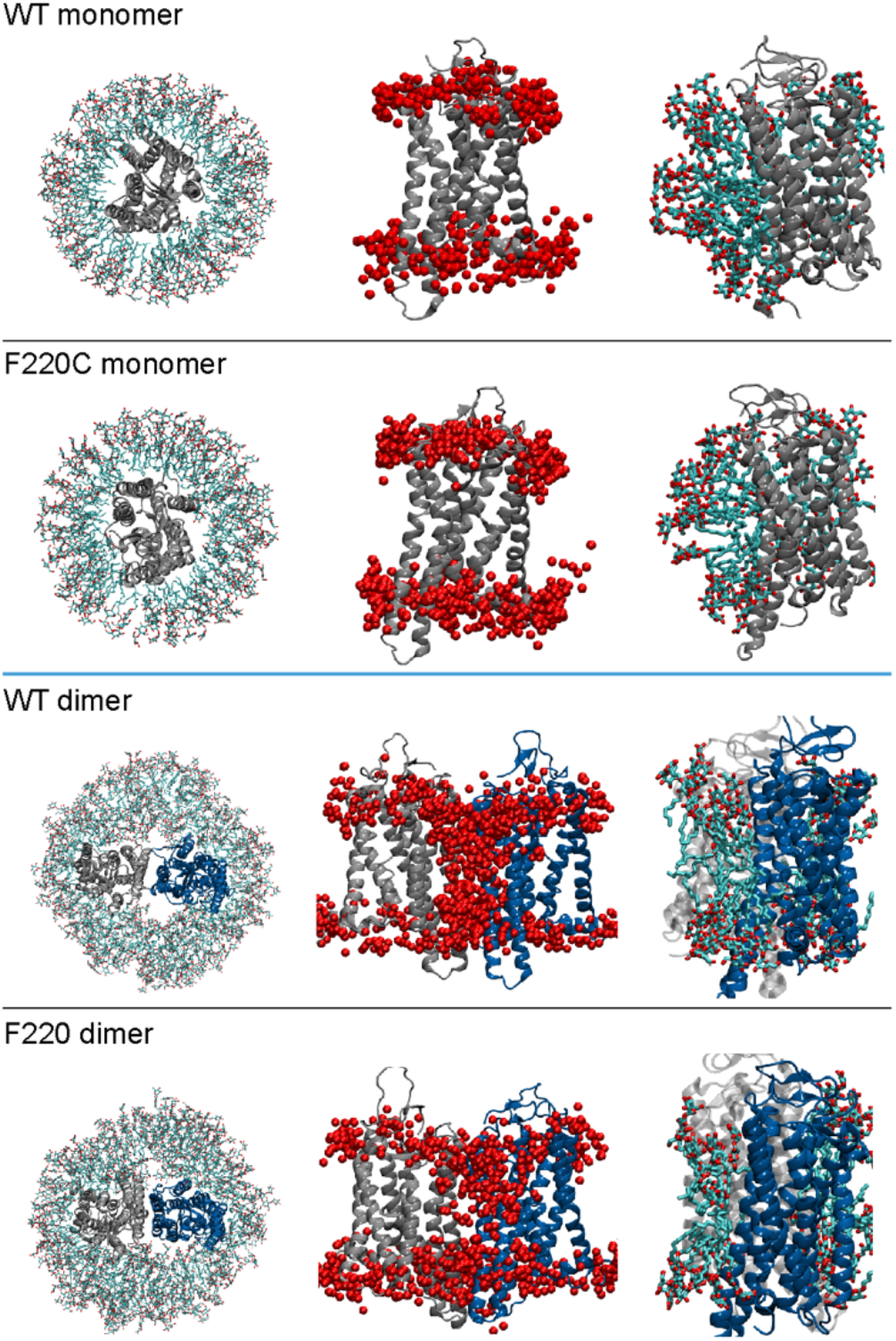
Structure of WT and F220C opsin monomers and dimers revealed by MD simulation. WT and F220C opsin monomers (top two rows) and dimers (bottom two rows) in DDM micelles. The snapshots on the left side of all rows show initial conformations of the respective systems: detergent molecules are shown as sticks, the protein is shown with its extracellular side on top (the monomer protein (tope two rows) is in grey, and the two protomers of the dimer constructs (bottom two rows) are in grey and blue). The middle panels show a side-view overlaid with locations of the O5 oxygen atom of the DDM headgroup (red spheres) that were within 4Å of the protein at least in one frame of the respective trajectory. The right panels show close up views of the protein constructs highlighting arrangement of the detergent molecules around TM5 and TM6 helices.

To calculate RHM, we used a computational approach based on quantifying the surface area involved in unfavorable hydrophobic-hydrophilic interactions ^24–26^. Figure S1 shows residues on TM5 and TM6 helices with high RHM values (> 1k_B_T) in simulations of the WT and F220C opsin monomers and dimers in micelles (the RHM values for the other parts of the protein were similar for all the systems). Overall RHM energy (F) stemming from these residues in the WT and F220C monomer systems (F_MONO_) is 6.4 ± 0.6 k_B_T and 6.2 ± 0.4 k_B_T, respectively. For the dimer systems, the RHM values (F_DIMER_) are significantly higher, 32.8 ± 3.5 k_B_T and 29.7 ± 3.4 k_B_T, for the WT and F220C dimers, respectively. Defining the RHM contribution to the dimerization energy as F_DIMER_- 2*F_MONO_, we find that the difference in RHM for the residues in TM5 and TM6 between the WT systems is 20 k_B_T and between the F220C systems is 17 k_B_T, suggesting that dimerization in a ‘high DDM’ environment is strongly unfavorable for both the WT and F220C systems. While our simulations consider only one of several possible modes of opsin dimerization (Table S1), these results are consistent with the experimental findings that both WT and F220C opsin proteins are monomeric at high detergent concentrations. Importantly, our results suggest that RHM may be a key factor in determining energy cost of dimerization during the *in vitro* reconstitution process as the detergent is gradually withdrawn from the system. Perhaps during detergent withdrawal, the energetic cost is significantly diminished for WT opsin, enabling it to self-associate. A detailed analysis of this process will be the subject of future work.

### Dimerization of both WT and F220C opsin in the plasma membrane revealed by ensemble FRET measurements

We next considered the self-association behavior of WT and F220C opsin in the membrane. We expressed N-terminal SNAP-tagged variants of WT and F220C opsin in HEK293T cells (Figure 3A). The proteins were labeled with cell-impermeant red and far-red SNAP dyes (SBG-JF_549_ and SBG-JF_646_) (28), thereby enabling analyses to be performed on the protein population expressed at the plasma membrane. A mixture of SNAP fluorophores was used to generate donor and acceptor labeled proteins in a 1:2 ratio. Quantification of fluorescence from several fields of cells revealed that all constructs were expressed at a comparable level (Figure 3B). FRET, quantified by measuring the increase in donor fluorescence upon bleaching of the acceptor with brief exposure to high intensity laser illumination, was observed at a similar level for both constructs indicating a degree of self-association (Figure 3C). These results indicate that both opsins dimerize/multimerize similarly in the plasma membrane, consistent with the result reported by Mallory et al. (17).

**Figure 3.**
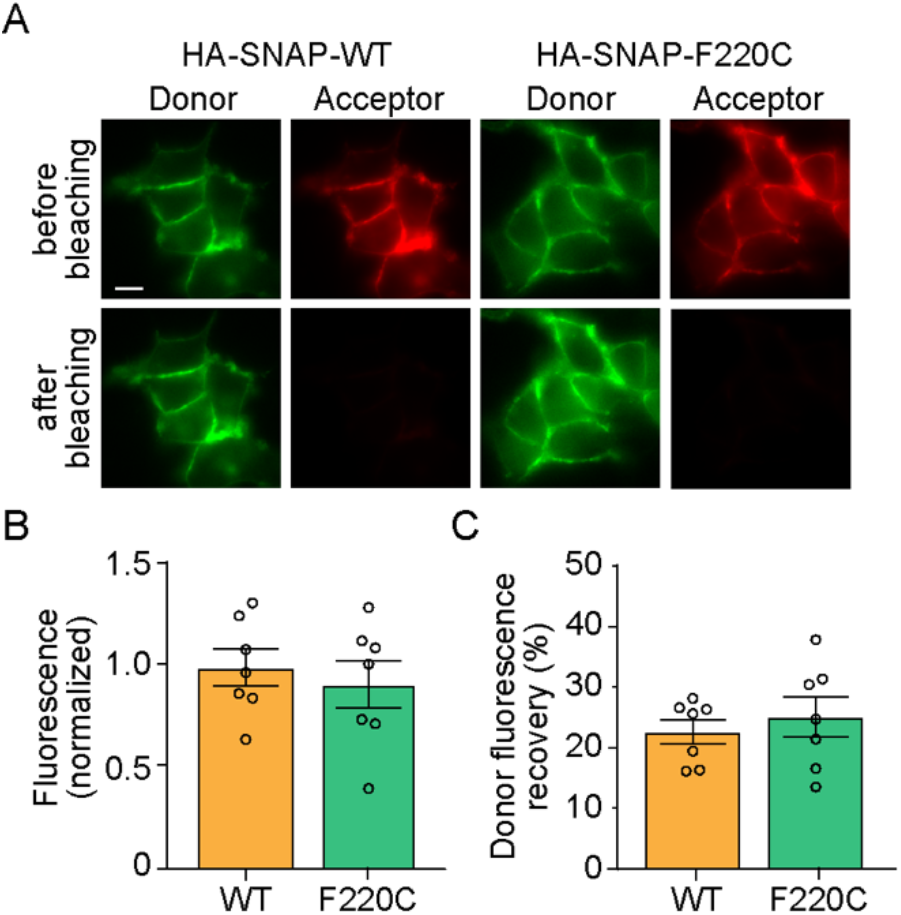
Dimerization of WT and F220C opsins in the plasma membrane of HEK293 cells. **(A)** Fluorescence images of HEK293T cells expressing HA-SNAP-WT opsin or HA-SNAP-F220C opsin and labeled with SNAP donor (SBG-JF_549_) and acceptor (SBG-JF_646_) dyes. The top and bottom rows show images of cells taken before and after bleaching of the acceptor by high intensity 640 nm illumination. Scale bar, 10 μm. **(B)** Summary of acceptor fluorescence as a reporter of comparable surface expression for both constructs. **(C)** Summary of donor fluorescence recovery upon acceptor bleaching.

### Workflow of computational experiments to investigate the behavior of opsin in the membrane

Thus far, we have shown that WT and F220C opsin differ dramatically in their dimerization propensity as they reconstitute into membranes from a micellar environment (16) (Figure 1E-G), yet self-associate to comparable levels in a membrane environment based on FRET measurements (Figure 3C). Because of the distinct behaviors of WT and F220C opsins in micelles, we considered the possibility that the dimers formed by these proteins in the membrane may be structurally different. To test this hypothesis, we used computational experiments in which we applied a multiscale approach in the spirit of the DAFT (Docking Assay For Transmembrane components) methodology ^(29,30)^. In these experiments, large numbers of pairs of monomeric proteins (WT and F220C) are allowed to aggregate spontaneously in a lipid membrane from initially well-separated states. The workflow that we used for each protein system is depicted in Figure 4A and involves the following steps: 1) Monomer protein models (WT and F220C) are equilibrated with atomistic MD simulations; 2) The atomistic structures are transformed into Martini coarse-grained (CG) representations; 3) Two copies of the CG protein are embedded into a lipid bilayer in such a way that the distance between their center-of-mass positions is ~50Å (see Figure 4B) and the system is briefly equilibrated with Martini force-fields. For this stage, we considered seven starting conformations differing in mutual orientations of the two monomers in order to randomize the initial conditions of the system; 4) Extensive ensemble CG MD simulations of spontaneous aggregation of the protein pairs are conducted in which every initial condition is simulated in 5 independent replicates, 120 μs each (~4.2 ms cumulative time in 35 replicates per construct); 5) The CG simulation trajectories are analyzed to identify preferred dimerization interfaces; 6) The selected dimer models are transformed into all-atom representations, in order to assess their stability and to quantify the energetics of dimerization with refined atomistic MD simulations. In the following, findings from these steps are presented in detail.

**Figure 4.**
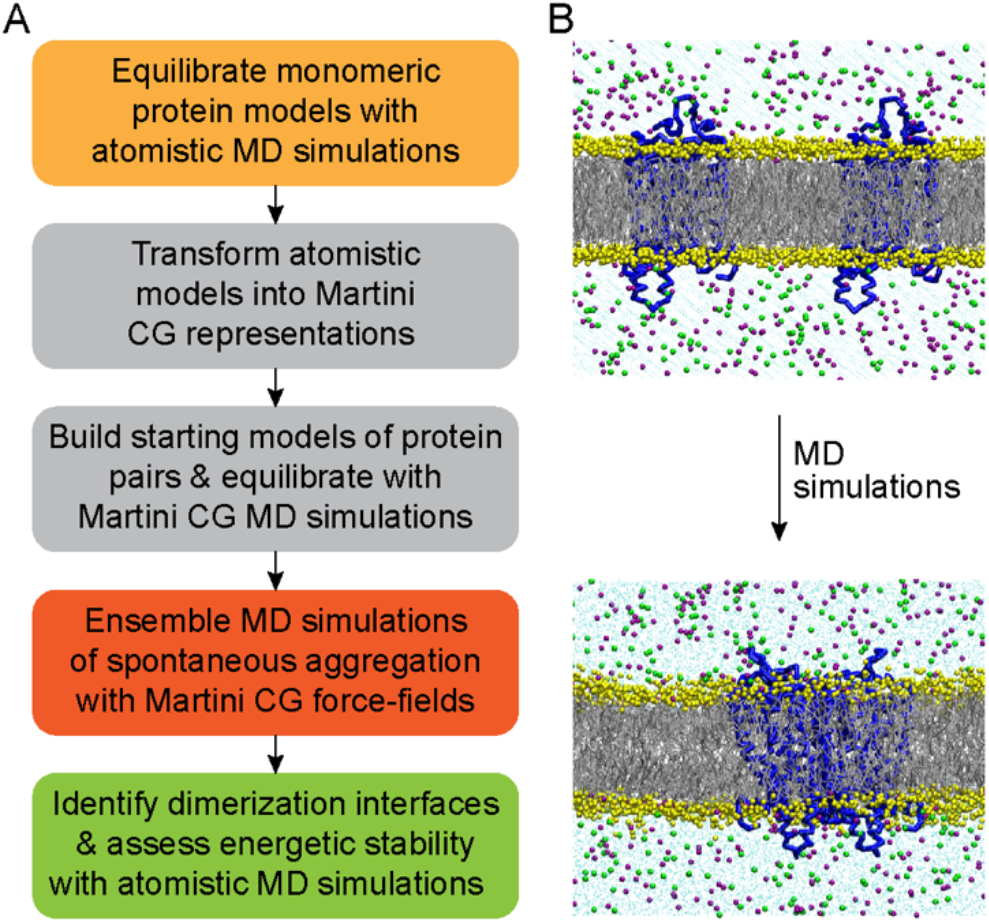
Set-up for computational analysis of opsin dimerization in a lipid membrane. **(A)** Multiscale computational protocol to investigate dimerization of WT and F220C opsins in a lipid membrane. **(B)** Starting (*top*) and final (*bottom*) snapshots of the WT opsin system after CG MD simulations. Initially well-separated monomers (center-of-mass distance of 50 Å) spontaneously aggregate on the 120 μs timescale of the CG simulations. In the snapshots, opsins are shown in blue cartoon, lipid phosphate beads in yellow spheres, and lipid tails in grey lines. Also depicted are salt ions in solution (green and purple spheres) and water beads (cyan dots).

### WT and F220C opsin monomers spontaneously aggregate into dimers during CG MD simulations

On the extensive timescales of the CG MD simulations, we observed many instances of spontaneous aggregation of opsin monomers (WT or F220C) into dimers. Indeed, as shown in Figures S2 and S3, out of 35 independent replicates of the WT system, dimerization was observed in 33 trajectories, whereas for the F220C construct, dimerization occurred in 30 out of 35 trajectories. Once formed, the dimer complexes remained stable for the remainder of the simulations. More detailed analysis of inter-protein interactions within the formed dimers revealed various modes of association involving different helices, and both symmetric and asymmetric dimer interfaces (Tables S1 and S2). To learn about the impact of the F220C mutation we compared dimerization interfaces formed by the TM5 helix, centering our attention on those trajectories that resulted in symmetric interfaces mediated by TM5-TM5 interactions (6 trajectories for WT system and 7 – for the F220C, see Tables S1 and S2).

### CG MD simulations identify two modes of dimerization along the TM5 helix in WT opsin

Figure S4 shows the frequency of pair-wise interactions for TM5 residues in the six CG trajectories of WT opsin in which dimerization was achieved through TM5-TM5 interactions. The data reveal two distinct modes of dimerization. One mode of dimerization (Mode 1), observed in trajectories 13 and 21 (Figure 5A and Figure S4), features extensive interactions along the intracellular end of TM5, involving residue stretch 204^5.39^-228^5.63^ (superscripts refer to residue labeling according to the Ballesteros-Weinstein generic residue numbering scheme for GPCRs (31)). The most notable interactions stabilizing this mode are found between pairs of F221^5.56^ and F228^5.63^ residues from opposing monomers. This dimerization mode includes additional stabilizing interactions along the extracellular part of TM6, as seen in Figure 5A (also Figure 5D, *left column*). In the second mode of dimerization (Mode 2), observed in trajectories 0, 27, and 31 (Figure 5A and Figure S4), dimers are mostly stabilized by pair-wise interactions between a shorter stretch of residues at the intracellular end of TM5, 228^5.63^-232^5.67^, as well as interactions between residues 220^5.55^ from the two protomers of the dimer. This mode of association is further stabilized by contacts along the extracellular side of TM6 helix (region between 273^6.56^ and 278^6.61^, see Figure 5A and also Figure 5E, *left column*). Lastly, trajectory 7 (Figure S4) shows relatively weak interfacial interactions involving 208^5.43^-212^5.47^ residue stretch in TM5.

**Figure 5:**
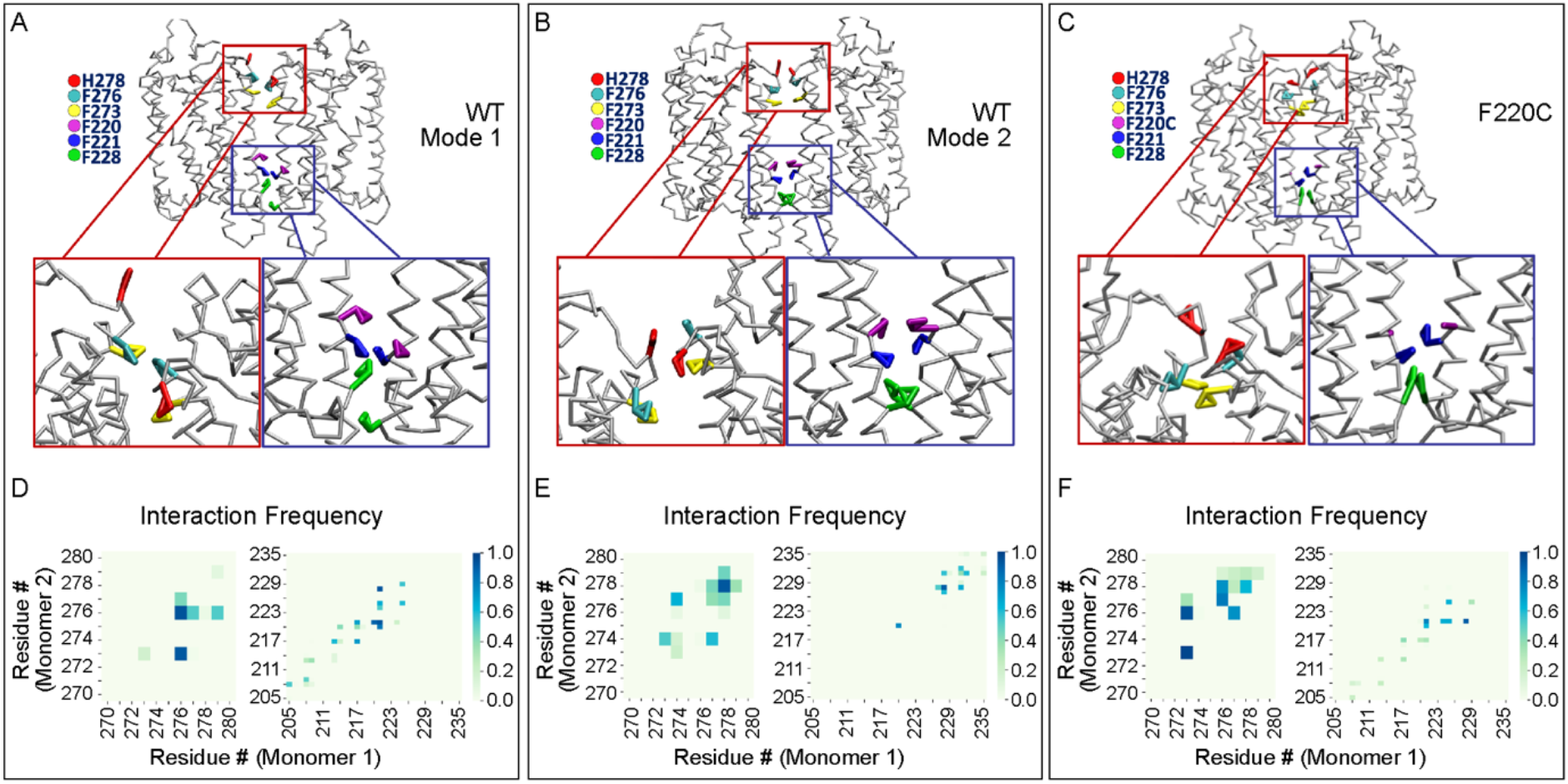
Modes of dimerization of the WT and F220C opsin. (**A-C**) Structural snapshots corresponding to the modes of dimerization of the WT (Modes 1 and 2 shown in panels A and B respectively) and F220C (C) opsin constructs from the CG MD simulations. Insets in red and blue rectangles show magnified views of interactions between pairs of TM6 and TM5 residues, respectively. Key residues participating in the interactions (H278^6.61^, F276^6.59^, F273^6.56^ in TM6 and F220^5.55^, F221^5.56^, F228^5.63^ in TM5) are depicted in different colors and are labeled. (**E**) Pairwise contact frequencies between residues on TM5 (*right columns*) and TM6 (*left columns*) in the dimerization modes shown in panels A-C respectively.

### F220C opsin shows a single mode of dimerization along the TM5 helix which is similar to one of the modes identified for the WT protein

Figure S5 shows the frequency of pair-wise interactions between TM5 residues for the six CG trajectories of the F220C opsin in which dimerization was achieved via TM5-TM5 interactions. The data reveal that all six trajectories evolve to a common dimerization interface in which inter-monomer contacts are stabilized through the 204^5.39^-228^5.63^ residue stretch in TM5. This dimerization interface is similar to that seen in Mode 1 dimerization in the WT system (compare Figure 5A and C, also Figure 5D and F).

### All-atom MD simulations structurally refine dimerization interfaces from the CG studies and reveal that F220C opsin forms energetically more stable dimers compared to the WT protein

The analysis of the CG MD simulations described above revealed the overall structural characteristics of the WT and F220C opsin dimers. In order to refine these structural features at the atomistic level and quantitatively compare the stability of the dimerization interfaces seen in the different constructs, we performed all-atom MD simulations. To this end, we selected representative structures from the CG simulations corresponding to the dimer arrangements described above (i.e. Mode 1 and 2 dimers for WT opsin, and a single dimer for F220C opsin, Figure 5A) and converted these CG dimers into all-atom representations. The all-atom structures were then subjected to ensemble MD simulations, in which the three dimer models were each run in 6 independent replicates (900 ns cumulative time per dimer construct). The protein constructs were structurally stable in these atomistic simulations as evinced by relatively small values of root-mean-square deviation (RMSD) of backbone atoms (Figure S6). Analysis of the frequency of pairwise interactions of the all-atom simulations showed largely the same trend as the CG simulations. Namely, as shown in Figure S7, both the Mode 1 WT dimer and the F220C dimer were stabilized by interactions in the 204^5.39^-228^5.63^ segment of TM5, whereas the Mode 2 WT dimer was maintained by interactions in the 228^5.63^-232^5.67^ region of the TM5 helix (although interactions between F220C residues on the two protomers were no longer observed in the all-atom simulations).

Importantly, quantification of dimerization energies revealed that F220C opsin formed energetically more stable dimers than the WT protein. Thus, we performed Molecular Mechanics - Generalized Born Surface Area (MM-GBSA) analysis of the all-atom trajectories to calculate the strength of dimerization for each system, taking into account the energetic cost of de-solvating the binding interface from its membrane and aqueous environment. In Figure 6, MM-GBSA binding energies are plotted separately for each all-atom trajectory. The data reveal significantly stronger binding of the F220C dimer compared to the WT dimers (compare WT1 and WT2 with F220C, see also Table S3). Analysis of sidechain and backbone atom contributions to the overall binding energetics (Figure S8) revealed that both contribute to the stability of the F220C dimers, suggesting that the energy difference between the WT and F220C dimers is the result not only of local sidechain rearrangements but also larger-scale conformational changes. To identify these changes, we next analyzed the structural differences in the WT and F220C dimers. In particular, we investigated the mutual arrangement of the two protomers in the dimers.

**Figure 6:**
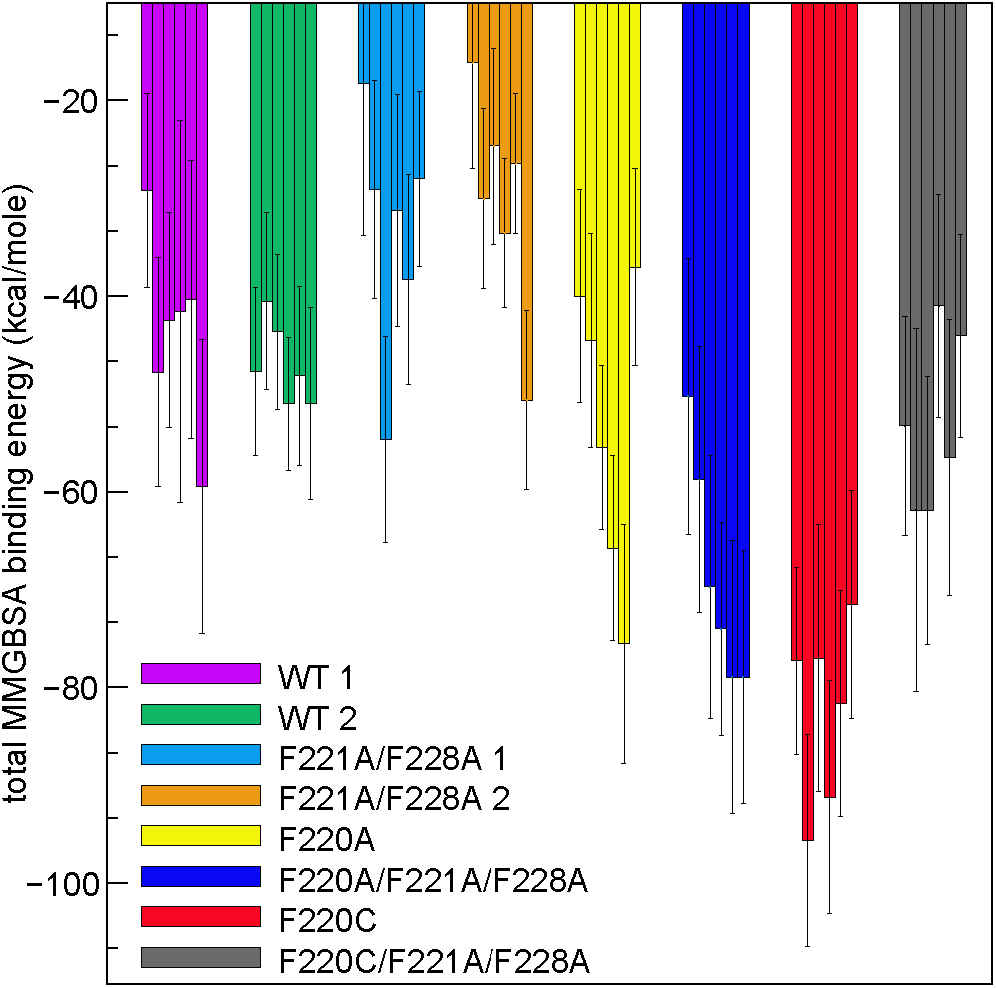
Binding energies of dimerization, quantified with the MMGBSA approach, for different opsin constructs. Data for six independent replicates per construct are shown separately. See also Table S3.

### Two protomers of the F220C opsin dimers orient in an unusually splayed conformation promoting widening of the intracellular vestibule and concomitant increase in hydration of the protein interior

Our analysis revealed that the F220C opsin dimer has a propensity to adopt conformations in which the TM5/TM6 helices from the two protomers are strongly tilted with respect to each other (Figure 7A, B). To quantify this mode of interaction, we defined the splay angle (α) between vectors representing the orientation of the TM5 helix in each protomer (each vector connects Cα atoms of residues 200 and 234). We constructed histograms of α values sampled along the trajectories. As shown in Figure 7C, the WT (Mode 1 and Mode 2) and F220C dimers sample conformations characterized by different splay angles. Whereas the dimers of WT opsin mostly sample lower splay angles (α<40°), F220C dimers sample conformations with larger α angles, with the splay angle reaching values around 50° and even higher. The time-evolution of the splay angle in the all-atom trajectories (Figure S9, see WT1, WT2, and F220C) revealed that the higher value for the splay angle in the F220C dimer was already present in the initial stages of the trajectories, suggesting that the splayed conformation developed dynamically in the CG simulations and remained stable on the timescales of the all-atom trajectories. To verify this, we calculated the α angle in CG trajectories of the F220C system and compared its behavior before and after the dimer formation event in these simulations. As shown in Figure S10A-B, α samples a wide range of values when the two opsin proteins are diffusing as monomers. But once the dimer is formed, α becomes confined to a narrow range of large values (Figure S10C), suggesting that the splaying in F220C is due to dimerization.

**Figure 7:**
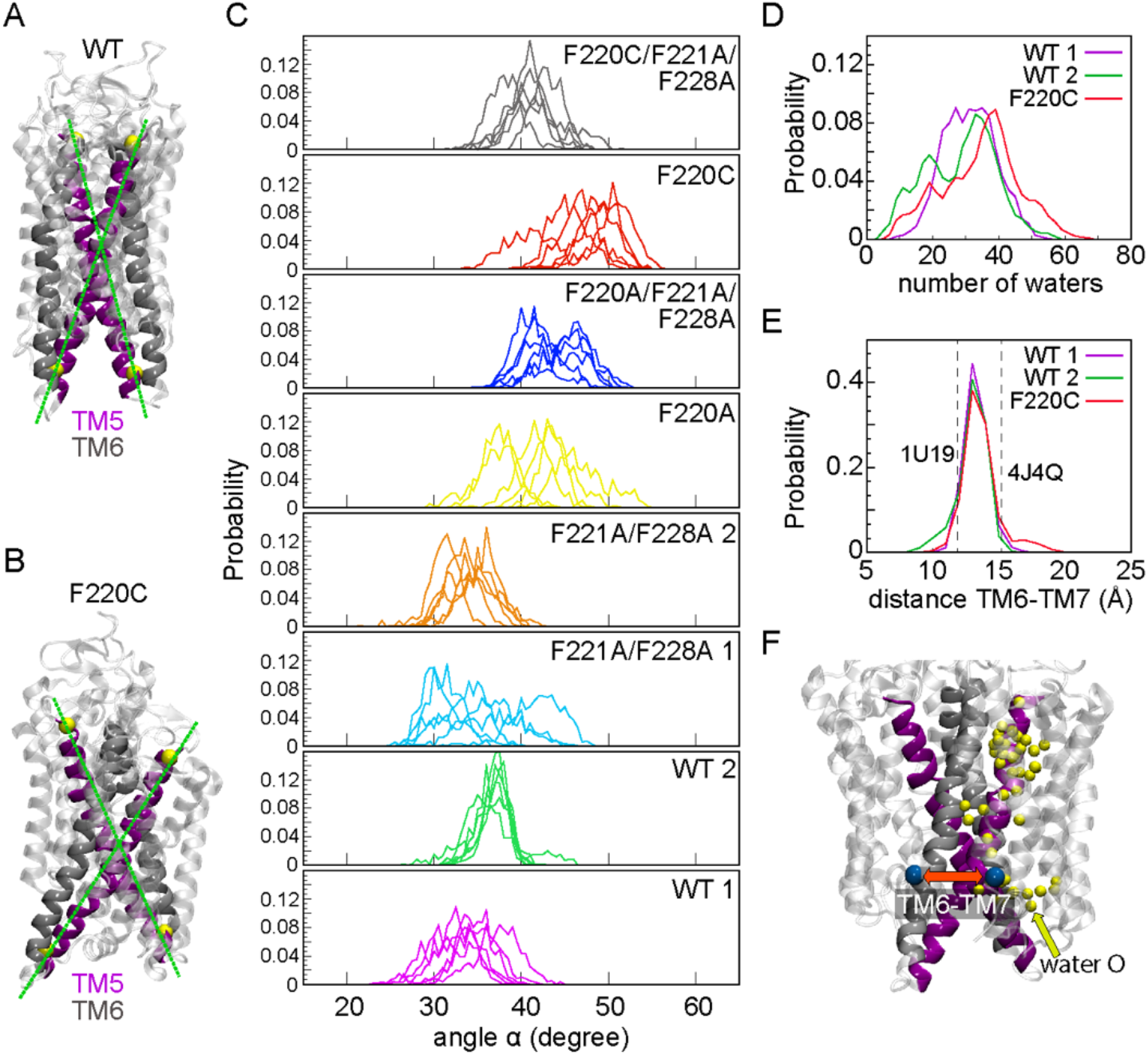
(**A-B**) Snapshots from all-atom MD simulations comparing mutual orientations of the two monomers in dimers of the WT (A) and F220C (B) constructs. TM5 and TM6 helices are shown in purple and silver, respectively. The rest of the protein is in transparent white color. Solid lines connect Cα atoms of residues 200 and 234 (yellow spheres) used to define the orientation of TM5 helix in each monomer (see text). (**C**) Probability distributions of the splay angle a between the two TM5 helices in the dimers of various constructs described in the text. The color coding of the systems follows the same pattern as in Figure 6. The results for each replicate are shown separately (i.e. 6 plots per constructs). (**D**) Histogram of water counts in the interior of each monomer of the WT (Modes 1 and 2) and F220C dimers. For this analysis, in each trajectory, for a given dimer, water oxygen atoms within 7A of the sidechains of residues 264, 268, 272, 127, and 131 were counted in each monomer. The data was then combined for two monomers and for all the trajectories per construct (i.e. WT 1, WT 2, and F220C) and binned using a bin size of 2. (**E**) Histogram of C_α_-C_α_ distance between residues 252 and 308 on each monomer of the WT (Modes 1 and 2) and F220C dimers. To construct the histogram, the data for each monomer and for all trajectories of each construct was combined and binned using a bin size of 1Å. (**F**) A snapshot of the F220C dimer as in panel B, highlighting water penetration (yellow spheres show water oxygens in the protein interior) and the distance between TM6-TM7 distance reported in panel E (blue spheres are Cα atoms of residues 252 and 308).

The consequences of the observed criss-cross orientation of the monomers in the F220C dimer are seen in the widening of the area between helices TM5/TM6 and TM7, and concomitant increase in the hydration of the protein interior. Indeed, as shown in Figure 7D, the distribution of the number of water molecules in the protein interior is shifted towards larger values in the F220C system compared to WT (Modes 1 and 2). At the same time, a small fraction of F220C dimers sample conformations in which the intracellular ends of TM6 and TM7 helices are separated by large distances, 17-20Å (see Figure 7E, also the snapshot in Figure 7F). The WT dimer is seen to mostly sample shorter TM6-TM7 distances, < 15Å. In this respect, it is important to note that the same distance measures ~15Å in the opsin model from PDBID 4J4U (GaCT peptide-stabilized structure) and ~11Å in the 1U19 model (retinal-bound, inactive form of the protein). Thus, our results suggest that the F220C mutation promotes widening of the intracellular vestibule in the corresponding opsin dimers, potentially enhancing the ability of the receptor to couple to effector proteins, such as transducin.

### The splayed conformation in the F220C dimer is a direct consequence of the mutation which alters residue interactions along the dimer interface

To identify the molecular mechanism underlying the observed splaying of the F220C dimer, we analyzed interfacial interactions in the WT and F220C dimer systems. The analysis revealed a subtle difference in the pattern of interfacial pairwise residue interactions in the two dimer systems. Thus, in the WT dimer (Mode 1), residue F221^5.56^ in one protomer is engaged in aromatic interactions with either F220^5.55^ or F228^5.63^ in the opposite protomer (Figure 8). In the F220C dimer, on the other hand, F221^5.56^ interacts mostly with just F228^5.63^ (Figure 8C) since position 220 is no longer occupied by a hydrophobic aromatic amino acid. The major role of the F221^5.56^ residue in stabilizing F220C dimers is exemplified by the larger contribution of this residue to the overall binding energy in the F220C system compared to the WT system as inferred from the per-residue decomposition of the MM-GBSA binding energies. Indeed, as shown in Figure S11, the side-chains of F221^5.56^, F228^5.63^, and I217^5.52^ are among those that contribute the most to the stability of the dimer interfaces of the WT (Mode 1) and F220C systems. Importantly, Figure S11 reveals that per-residue binding energy for F221^5.56^ is notably larger in the F220C dimer construct. Thus, in 5/6 trajectories of the F220C system, the top contribution to dimer stabilization stems from the sidechain of residue F221^5.56^, which contributes between −3.8 and −4.6 kcal/mole to the overall binding energy (Figure S11).

**Figure 8:**
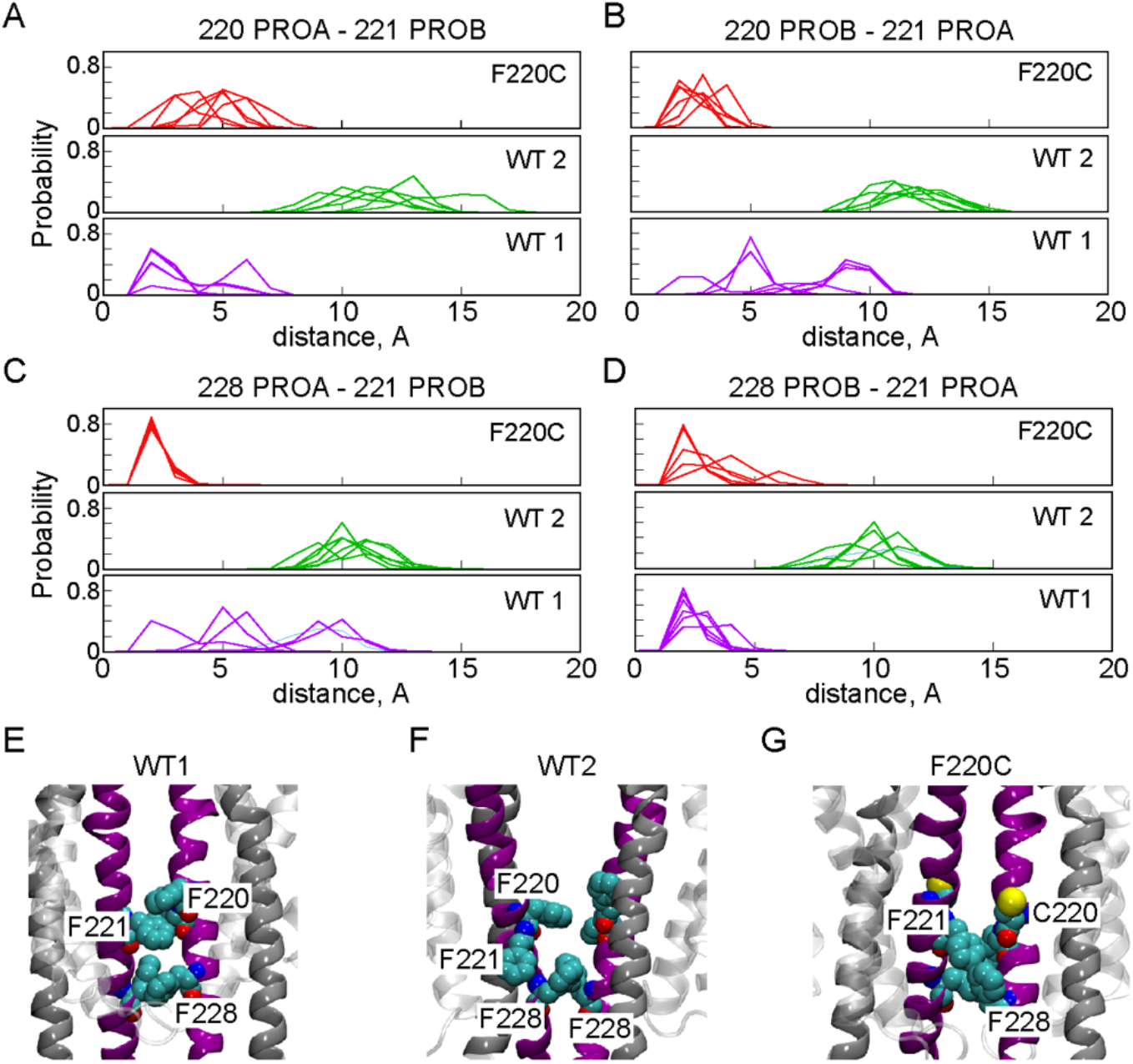
(**A-B**) Probability distribution of the minimum distance between residues F/C220^5.55^ and F221^5.55^ on the opposite monomers of dimers formed by WT opsin (Modes 1 and 2 - WT1, WT2) and F220C. (**C-D**) Probability distribution of the minimum distance between residues F228^5.53^ and F221^5.55^ on the opposite monomers of dimers formed by WT opsin (Modes 1 and 2 – WT1, WT2) and the F220C mutant. PROA (panels A, C) and PROB (panels B, D) denote two monomers of the dimer, and the results are shown separately for all six replicates for each construct. (**E-G**) Close-up view of molecular interactions along TM5 helix in the F220C dimer (G) and in the Mode 1 and Mode 2 WT dimers (E and F respectively). Relevant residues are shown in space-fill and labeled. TM5 and TM6 helices are shown in purple and silver, respectively.

Because F221^5.56^ is about two helical turns above F228^5.63^ on the opposite protomer, stable interactions between these residues in the F220C dimer requires the phenyl ring of F228^5.63^ to position “upward” (Figure 8G). To sustain such an orientation, the intracellular part of TM5 moves outward resulting in the overall tilt of this helix with respect to the other protomer. Thus, the splayed configuration of the F220C dimer appears to be a direct consequence of the mutation at the 220 position, which alters pair-wise residue interactions at the dimer interface.

To test whether this peculiar, splayed arrangement is specific to the F220C mutation, we computationally probed several other mutations in TM5. We considered an Ala substitution at the 220 position (F220A), and, given their importance in interfacial interactions (Figure 8A-D), we also introduced Phe-to-Ala mutations at positions 221 and 228 using Mode 1 and Mode 2 structures of the WT dimers to create F221A/F228A 1 and F221A/F228A 2 constructs. Lastly, we created the triple mutants F220C/F221A/F228A and F220A/F221/F228A. The new systems were subjected to all-atom MD simulations in the same manner as the F220C and the WT constructs (i.e. 6 replicates, 150 ns duration each).

Calculation of the α splay angle in the new mutant systems revealed that all the constructs that had a mutation at position 220 were, on average, characterized by larger splay angle compared to the ones in which this site was not altered (Figures 7C and S9). Interestingly, F220C still showed the largest splay angle among all the systems. MM-GBSA calculations (Figure 6) revealed that F221A/F228A 1 and F221A/F228A 2 constructs had WT-like binding energies which were significantly lower compared to the binding energies for the F220C system (see Table S3). On the other hand, all the constructs in which position 220 was mutated were characterized by binding energies more similar to that found for F220C compared with constructs in which the 220 position was unaltered. These results suggest that a side chain at position 220 position is crucial for the stability of the TM5-TM5 dimerization interface.

To test experimentally the effects of the mutations at 220, 221, and 228 positions, we constructed single, double and triple mutants of opsin (F1 (F228A), F2 (F221A, F228A), and F3 (F220C, F221A, F228A)), expressed them along with WT protein in HEK293 GnTI^-/-^ cells and purified them by affinity chromatography (Figure 9A). The yield of the constructs was similar, indicating that the various mutations do not have a detrimental effect on protein structure. We reconstituted the proteins into large unilamellar vesicles and used our end-point scramblase assay to measure the fraction of vesicles able to scramble lipids as a function of the amount of protein that was reconstituted into a fixed number of liposomes. The data clearly show that F3 reconstitutes as a monomer, whereas F1 and F2 opsins that lack the F220C mutation reconstitute as dimers, similar to WT protein (Figure 9B).

**Figure 9:**
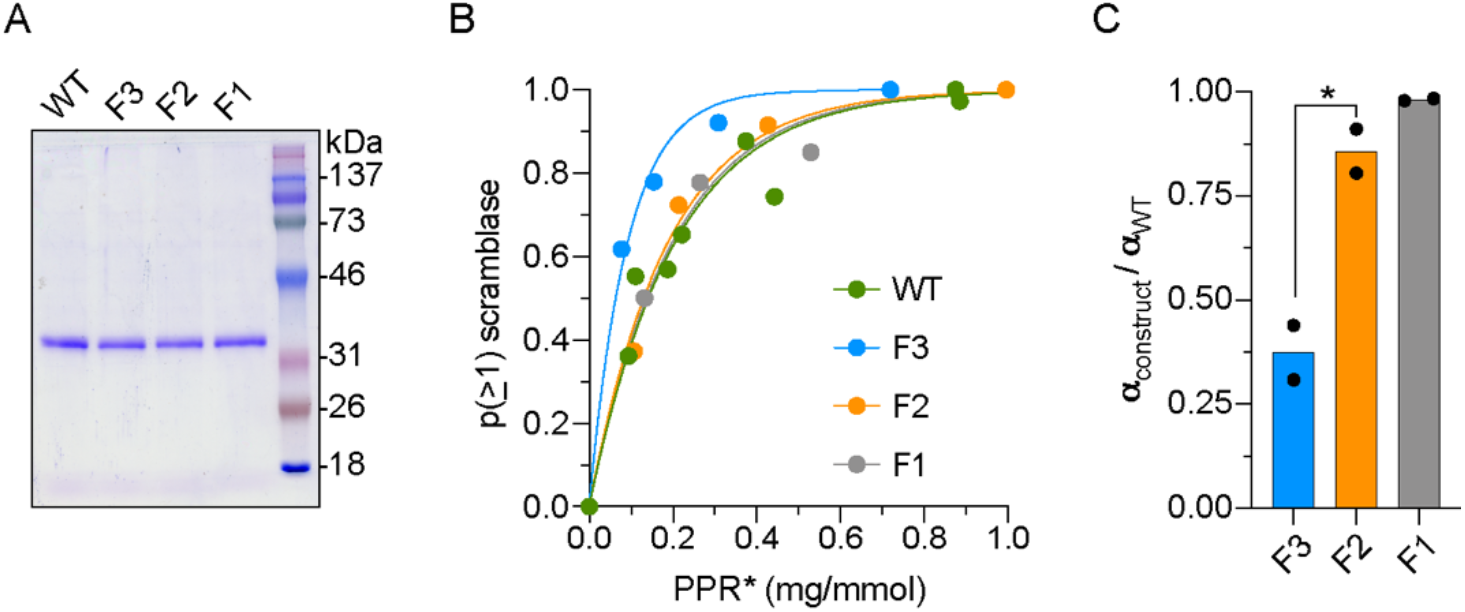
Reconstitution of WT opsin and TM5 mutants into liposomes. **(A)** Coomassie-stained SDS-PAGE of purified WT opsin and the F1 (F228A), F2 (F221A, F228A) and F3 (F220C, F221A, F228A) opsin constructs. Molecular weight markers are indicated. **(B)** Protein dependence of the reconstitution of vesicles with scramblase activity. NBD-PC-containing vesicles were reconstituted with different amounts of the indicated opsin construct and the extent of fluorescence reduction on dithionite treatment was determined by curve-fitting. The data were processed to calculate the probability “p(≥1) scramblase” that an individual vesicle in the ensemble possesses a functional scramblase. The graph shows “p(≥1) scramblase” versus the corrected protein/phospholipid ratio (PPR*) of the sample (the correction eliminates the contribution of vesicles that are refractory to reconstitution). **(C)** Ratio of mono-exponential fit constants (a) obtained from analyses of the reconstitution of WT and mutant opsins similar to (and including) the analysis shown in Panel (B). The ratio α_construct_/α_WT_ ~1 for F1 and F2 opsins, indicating that these proteins reconstitute similarly to WT opsin as dimers or higher order multimers. However, α_F3_/α_WT_ ~0.4, consistent with reconstitution of F3 as a monomer (α_WT_ = 0.25 ± 0.07, mean ± S.D.). The bar chart is constructed from the results of three independent experiments, each involving 4-5 reconstitutions at different protein/phospholipid ratios (*, p =0.029, unpaired t-test (two-tail)).

As an additional test of specificity of the splayed arrangement for the F220C substitution, we considered the splayed conformation of the F220C dimer (α ~48°), and introduced in this construct the wild type Cys residue into position 220 (C220F mutant). We then initiated 6 independent MD simulations of this dimer system. As shown in Figure S12, during ~300 ns runs (excluding the standard equilibration protocol described in Methods), in 3/6 trajectories α started to decrease from its initial high value, suggesting that the splayed arrangement indeed is stabilized by the F220C mutation.

### Conformational differences in WT and F220C opsin dimers revealed by ensemble FRET experiments using constructs with a C-terminal SNAP-tag

The MD simulations described above indicate that the conformational pose of the F220C dimer differs from that of the WT dimer. We considered the possibility that this difference, not evident in the FRET experiments with N-terminal SNAP-tagged constructs (presented in Figure 3), might be revealed if the SNAP tag were located at the C-terminus. Accordingly, we expressed C-terminally SNAP-tagged WT and F220C opsins in HEK293 cells, labeled the tags using membrane-permeant donor and acceptor fluorophores, and assessed FRET (Figure 10). The constructs were expressed at a similar level (Figure 10B), but now the extent of donor recovery was significantly greater for F220C opsin compared with WT (Figure 10C). Importantly, the same fluorophores were used for both N- and C-terminal SNAP-tagged constructs, eliminating the possibility that differences are due to photophysical differences. While this result does not permit a detailed structural comparison of WT and F220C dimers, it clearly shows that the SNAP tags are positioned differently in the F220C dimer compared with WT, consistent with the prediction of the MD simulation that the F220C mutation stabilizes the intracellular end of the protein juxtaposed to the C-terminus in a different conformation compared to the WT protein.

**Figure 10:**
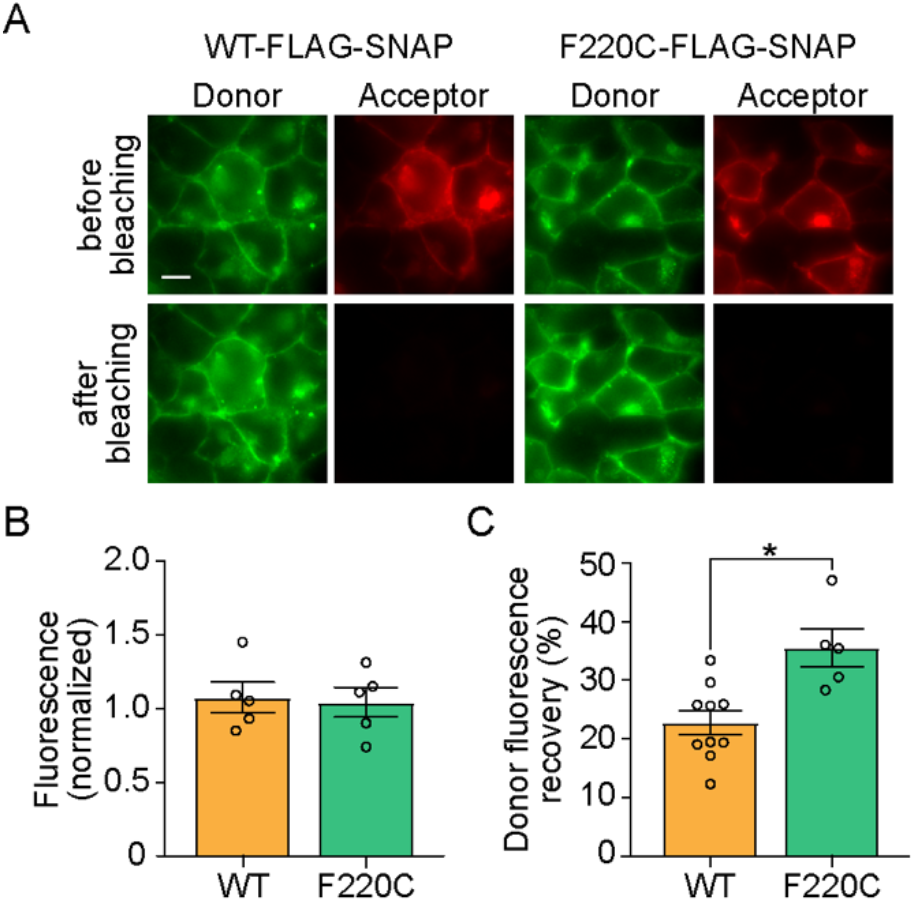
Distinct dimer conformations of WT and F220C opsins in the plasma membrane of HEK293 cells. **(A)** Fluorescence images of HEK293 cells expressing WT-FLAG-SNAP opsin and F220C-FLAG-SNAP opsin labeled with SNAP donor (BG-JF_549_) and acceptor (BG-JF_646_) dyes. The top and bottom rows show images of cells taken before and after bleaching of the acceptor by high intensity 640 nm illumination. Scale bar, 10 μm. **(B)** Summary of acceptor fluorescence as a reporter of comparable surface expression for both constructs. **(C)** Summary of donor fluorescence recovery upon acceptor bleaching, indicative of FRET (*, p =0.013, unpaired t-test (two-tail)).

### Conclusion

The F220C rhodopsin mutant was originally discovered in a study of 88 patients/families with a history of RP (32). Based on *in vitro* reconstitution studies we originally proposed (16) that F220C differed from WT opsin in its ability to dimerize *en route* to reconstitution into lipid vesicles. We suggested that this deficiency in an otherwise normal rhodopsin could account for its disease association. However, subsequent work (17) unexpectedly indicated that F220C opsin could dimerize in the membrane, and recent studies of the F220C-knockin mouse model revealed only minor phenotypic consequences of this mutation (18). We now present an expansive characterization of this enigmatic opsin mutant, using a combination of experimental and *in silico* studies. Using new, fluorescence-based experimental approaches we show that F220C opsin, like WT opsin, is a monomer when expressed in HEK293 cells and isolated using 1% (w/v) DDM. However, reduction of detergent levels causes the WT protein to dimerize whereas F220C remains monomeric. Remarkably, MD simulations showed that the micellar environment is energetically unsuited to opsin dimers, raising the question of why the WT protein dimerizes at all under these conditions. Further work will be needed to understand this phenomenon. The behavior of the F220C mutant in the membrane also contrasted with that of the WT protein. Whereas we could show that F220C dimerizes in the membrane as previously reported, MD simulations revealed that it forms an energetically favored dimer when compared with the WT protein. The conformation of the F220C dimer is unique, with the TM5/TM6 helices tilted to form a splayed, criss-crossed structure promoting widening of the intracellular vestibule of each monomer in the dimer. This could potentially enhance the ability of the receptor to couple to effector proteins, such as transducin, and also influence the higher order assembly of rhodopsins in disc membranes. While it is intriguing that the difference in quaternary structure between WT and F220C opsins does not immediately translate into a more dramatic phenotypic consequence in the F220C-knockin mouse model, the unusual nature of the F220C dimer may account, at least in part, for the observed phenotype and disease association in humans.

## Experimental Procedures

### Computational Methods

#### Molecular Constructs for all-atom MD simulations of opsin monomer systems

All computations were based on the X-ray structure of retinal-free opsin (PDBID 4J4Q) (33). This structure also contains bound synthetic GaCT peptide which was not considered here. Using the CHARMM-GUI web interface (34), the wild type (WT) and F220C opsin models were prepared and embedded in an all-atom phospholipid bilayer containing a 9:1 mixture of POPC (1-palmitoyl-2-oleoyl-*sn*-glycero-3-phosphocholine) and POPG (1-palmitoyl-2-oleoyl-*sn*-glycero-3-phospho-(1’-rac-glycerol)) lipids, mimicking the composition of the reconstituted vesicles used for scrambling activity assays (3). The protein to lipid ratio was 1:362. After adding a solvation box containing 150 mM KCl the total system size was ~130,000 atoms.

#### All-atom MD simulations of opsin monomer systems

All-atom systems containing the WT and F220C mutant opsin monomers were subjected to ~130 ns MD simulations using NAMD 2.12 software and the latest all-atom CHARMM 36m force fields for protein, lipids and ions (35). The simulations followed the multi-step equilibration protocol prescribed by CHARMM-GUI before switching to unbiased runs. These simulations implemented the *all* option for rigid bonds, 2fs integration time-step, PME for electrostatics interactions (36), and were carried out in NPT ensemble under semi-isotropic pressure coupling conditions, at a temperature of 303 K. The Nose-Hoover Langevin piston algorithm (37) was used to control the target P = 1 atm pressure with the Langevin piston period set to 100 fs and decay time constant set to 50 fs. The van der Waals interactions were calculated applying a cutoff distance of 12 Å and switching the potential from 10 Å. In addition, the vdwforceswitching option was set to *on*.

#### Molecular constructs for coarse-grained (CG) Martini simulations

Using CHARMM-GUI, the opsin monomer models (WT and F220C) from the final frame of the respective all-atom MD trajectories were transformed into Martini-based coarse-grained (CG) structural representation (38,39). For each construct, a second copy of the CG protein was then created by translating the center-of-mass of the first one in *x/y* (membrane) plane by 50Å. The two CG opsins were randomly rotated with respect to each other and around *z*-axis to create 5 different mutual orientations of the pair of proteins. Using again CHARMM-GUI, these 5 opsin pairs were embedded into a 1099 lipid-size membrane containing 9:1 mixture of POPC and POPG lipids. The total system size, after solvating and ionizing steps was 38280 CG beads.

#### CG MD simulations of spontaneous aggregation of opsin monomers into dimers

For each protein construct (i.e. WT or F220C) and mutual orientation 7 independent CG MD simulations of 120 μs in length were performed using Gromacs version 5.1.4, resulting in 35 total simulations per construct with a cumulative time of 4.2 ms. The simulations implemented 20fs timestep, and were carried out in NPT ensemble under semi-isotropic pressure coupling conditions, at a temperature of 303 K. The Berendsen algorithm was used to control the target P = 1 atm pressure with the τp and *compressibility* parameters set to 5.0 ps and 3e-4 bar^-1^, respectively. The temperature was controlled by the v-rescale algorithm (separated for protein, membrane and solute).

#### Molecular Constructs for all-atom MD simulations of opsin dimer systems

Using CHARMM-GUI, selected opsin dimer CG models for the WT and F220C system (see Results) were transformed back into all-atom representation. For the WT system, two models were considered, termed, Mode 1 and Mode 2. For F220C, a single CG model was transformed. To create an all-atom model for the F221A/F228A double mutant system, Phe-to-Ala substitutions were introduced to Mode 1 and Mode 2 WT models, resulting in two starting conformations for the double mutant system, F221A/F228A 1 and F221A/F228A 2. The all-atom model for the F220A mutant was created by replacing Cys220 by Ala in the all-atom F220C dimer. Lastly, two triple mutant systems were created F220A/F221A/F228A and F220C/F221A/F228A by introducing F221A and F228A mutations into the F220A and F220C models. Together, 6 different dimer constructs were created, totaling 8 systems: WT (two starting models), F220C, F220A, F221A/F228A double mutant (two starting models), F220A/F221A/F228A, and F220C/F221A/F228A.

Using CHARMM-GUI, each of these 8 models were embedded in a 900 lipid-size membrane consisting of 9:1 mixture of POPC/POPG lipids. After adding a solvation box containing 150 mM KCl the total system size was ~320,000 atoms.

#### All-atom MD simulations of opsin dimer systems in lipid membranes

The eight all-atom dimer systems were subjected to ensemble MD simulations using NAMD 2.12 software (37). Each system was run in six 150ns long independent replicate, resulting in cumulative time of 900ns per construct. For these runs as well, the latest all-atom CHARMM 36m force fields for protein, lipids and ions, and the simulations used the same protocols and run parameters as described above for the all-atom monomer systems.

#### All-atom MD simulations of opsin dimer systems in detergent micelles

WT and F220C opsin monomers and dimers were simulated in micelles composed of DDM detergent. The initial structures of the dimer systems were taken from the all-atom MD simulations in lipid membranes in which the F220C dimer was in splayed conformation whereas in the WT dimer the two subunits were more aligned with respect to each other (see text). The same structures served as starting points for the monomer simulations. The protein constructs were inserted into DDM micelles, solvated and ionized using CHARMM-GUI. Protein-to-detergent ratio was 1:150 for the monomers and 1:300 for the dimers. Each system was first equilibrated using standard CHARMM-GUI equilibration protocol and then run with OpenMM 7.4 (40) in six independent replicates (WT systems – 150ns each, F220C monomer – 260ns each, F220C dimer – 300ns each). For these simulations as well, the latest all-atom CHARMM 36m force fields for protein, lipids and ions were used. The simulations used the same protocols and run parameters as described above for the allatom systems except 4fs integration timestep (with hydrogen mass repartitioning), and isotropic pressure coupling.

#### Calculation of dimerization energies from all-atom MD simulations using the MM-GBSA method

To quantify strength of opsin dimerization, we calculated binding energies using the MM-GBSA (Molecular Mechanics, Generalized Born Surface Area) (41). The approach combines molecular mechanics interaction energies from the CHARMM36 force field (42) with a generalized Born and surface area continuum solvation model to estimate solvation effects upon binding of macromolecules (in this case two monomers of the protein dimer). Protein conformations are sampled from all-atom MD trajectories, from which all water molecules, lipids and ions are discarded. Here we used the single-trajectory variant of the method, where unbound conformations of one protomer are obtained by discarding the atoms of the other protomer. Thus, the results do not take into account any reorganization energies associated with internal conformational changes upon binding. Despite this strong assumption, the approach has been widely used to compare strength of binding of different macromolecular complexes, with the useful possibility of performing approximate per-residue binding energy decompositions (43,44) that compare well to experimental alanine scanning (45–47).

MM-GBSA analysis was carried out on 25-150ns time-intervals of the all-atom trajectories using the implementation in the CHARMM software (48) complemented with in-house scripts. as described previously (49). Briefly, the dielectric constant inside the protein was set to 2 and we used the accurate Generalized Born Molecular Volume II (GBMVII) method (50) to calculate the polar solvation free energies. The non-polar solvation free energy was proportional to the solvent accessible surface area (SASA) with a coefficient of −0.0072 kcal/mol/Å^2^ (51). We used the Debeye-Hückel correction to account for ionic screening (52), with screening constant 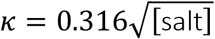 (expressed in Å^-1^) (53), where [salt] is the monovalent ion concentration taken to be 0.154 mol/l. The membrane was treated as a continuum slab with a dielectric constant varying from low in the membrane core to medium in the head-group region using the Heterogeneous Dielectric Generalized Born (HDGB) method (54). We used the improved dielectric profile and non-polar correction profile from Ref. (55).

Using this framework, to calculate binding energy in a specific dimer construct, MM-GBSA calculations were performed to obtain the solvation free energies separately for three molecular systems: for the dimer (G_DIMER_), and for the two monomers of the dimer separately (G_PROA_ and G_PROB_). Entropic contributions in the proteins were neglected, so that the dimerization free energy was then estimated from: ΔG_bind_=E_int_+G_DIMER_-(G_PROA_+G_PROB_), where E_int_ is the electrostatic and Van der Waals interaction energy from CHARMM36 with no distance cutoff. All terms in ΔG_bind_ being either local or pair-wise interactions, a decomposition is obtained by summing all local contribution and half of the pair-wise interactions for all atoms in a group (typically a residue or a side chain) (43,44).

#### Calculation of the RHM energies from all-atom MD simulations in micelle systems

RHM values for each protein residue was calculated as described previously (25–27,56). Briefly, the residual exposure of each residue (*SA_res,i_*) was quantified from Solvent Accessible Surface Areas (SASA) calculated from corresponding MD trajectories, as follows:

For hydrophobic residues, *SA_res,i_* is the surface area of the hydrophobic residues exposed to the micelle headgroup or water and is calculated as the SASA with the solute taken to be the protein and hydrophobic core of the detergent micelle (i.e. hydrocarbon tails of DDM molecules, atoms C1-C12 in CHARMM36m nomenclature).

For hydrophilic residues, *SA_res,i_* is the surface area of the residue embedded in the hydrophobic core of the micelle and is calculated as the difference of the SASA with the solute taken to be the protein only and the SASA with the solute taken to be the protein and the hydrophobic part of the micelle.

The corresponding energy penalty (*ΔG_res,i_*) can be approximated as linearly proportional to *SA_res,i_*: Δ*G_res,i_* = *σ_res_SA_res,i_*, with the constant of proportionality *σ_res_* taken to be 0.028 kcal/(mol. Å^2^) (57,58). For more details of the method please see Ref. (25–27,56).

### Experimental Methods

#### Materials

Anti-FLAG-M2 affinity agarose gel and 3X FLAG peptide were obtained from Sigma Aldrich, n-dodecyl β-D-maltoside (DDM) was from Anatrace, protease inhibitor cocktail Set V was from Calbiochem/EMD Millipore Corp, Transporter 5^™^ transfection reagent (PEI MAX) was from Polysciences Inc., Antibiotic-Antimycotic (100X, containing 10,000 units penicillin, 10,000 μg/ml streptomycin and 25 μg/ml amphotericin) was from Gibco, and spin columns (product number 69705) were from Pierce Biotechnology. HEK293S GnTI^-^ cells were a kind gift of the Meyerson laboratory (Weill Cornell Medical College). NEBuilder HiFi DNA Assembly Cloning Kit, Phusion polymerase, Dpn1 and T4 DNA ligase were from New England Biolabs, and primers were obtained from IDT. Phospholipids (1-palmitoyl-2-oleoyl-glycero-3-phosphocholine (POPC), 1-palmitoyl-2-oleoyl-*sn*-glycero-3-phospho-(1’-rac-glycerol)(POPG) and 1-palmitoyl-2-{6-[(7-nitro-2-1,3-benzoxadiazol-4-yl)amino]hexanoyl}-*sn*-glycero-3-phosphocholine (NBD-PC)) were obtained as stock solutions in chloroform from Avanti Polar Lipids. Bio-Beads SM2 resin was from Bio-Rad Laboratories. Sodium dithionite was from Sigma. SNAP-Surface 549 and SNAP-Surface 649 were from New England Biolabs.

#### Opsin constructs

We previously described a bovine opsin construct containing N2C and D282C mutations for enhanced thermostability, and a C-terminal tandem 3X-FLAG tag (16). The construct, here termed wild-type (WT), was prepared in the pMT3 expression vector. The F220C construct, derived from WT, was also previously described (16). Additional mutations were introduced into the WT construct to generate F1 (F228A) and F2 (F221A, F228A), and into F220C to generate F3 (F220C, F221A, F228A). Site-directed mutagenesis was achieved using the following primers: for F2, forward primer - **<u>GCC</u>** TGC TAT GGC CAG CTG GTG **<u>GCC</u>** ACC GTC AAG GAG GCT GCA GCC C, reverse primer - GAA GAT GAC AAT CAG CGG GAT GAT G, and for F3, forward primer - **<u>GCC</u>** TGC TAT GGC CAG CTG GTG **<u>GCC</u>** ACC GTC AAG GAG GCT GCA GCC C, reverse primer - GCA GAT GAC AAT CAG CGG GAT GAT G, and for F228A, forward primer - **<u>GCC</u>** ACC GTC AAG GAG GCT GCA GCC C, and reverse primer - CAC CAG CTG GCC ATA GCA GAA GA. Polymerase chain reactions (PCR) were performed using High Fidelity Phusion polymerase. PCR-amplified products (verified by electrophoretic separation using a 1% agarose gel) were treated with the Dpn1 restriction enzyme to remove the parent plasmid before being subjected to ligation (using T4 DNA ligase) and transformed into *Escherichia coli* DH5α cells. All constructs were sequenced to confirm the desired mutations.

N-terminally tagged HA-SNAP-Opsin (WT and F220C) clones were made by replacing the mGluR2 gene from a previously described N-terminal HA-SNAP-mGluR construct (59) with thermostable opsin-FLAG, using the NEBuilder HiFi DNA Assembly Cloning Kit. The HA-SNAP containing vector backbone was amplified using the forward and reverse primers TGAAGATCCCACACTCCTGCCC and ACGCGTGCCCAGCCCAGG, respectively, and opsin genes (WT and F220C) were amplified from the corresponding pMT3-opsin constructs (described above) using forward (<u>AGCCTGGGCTGGGCACGCGT</u>TGCGGTACCGAAGGCCCAAAC) and reverse (<u>GCAGGAGTGTGGGATCTTCA</u>CTTGTCATCGTCATCCTTGTAGTCG) primers containing the overlapping region (underlined) from the HA-SNAP-mGluR construct. The constructs were confirmed by sequencing.

#### Cell culture and transfection

Homogeneously *N*-glycosylated FLAG-tagged opsins were produced by transfection of HEK293S GnTI^-^ cells using the Transporter 5^™^ transfection reagent (PEI MAX). Briefly, cells were harvested from a confluent (80-90%) 10-cm plate cultured at 37°C in a 5% CO2 atmosphere in Dulbecco’s Modified Eagle Medium (DMEM) containing 10% fetal bovine serum (FBS) and antibiotic-antimycotic, and split into four 10-cm plates using the same medium. After 24 h when the cells were 60-70% confluent, the culture medium was replaced with 10-mL (per plate) fresh DMEM containing 4% FBS. The cells were incubated for a further 2 h before transfection. For each plate, a transfection mix was prepared by combining 1 ml 150 mM NaCl, plasmid DNA (20 μg of a 4:1 (w/w) mixture of the pMT3 plasmid carrying the opsin gene and pRSVTag carrying a gene encoding SV40 large tumour (T) antigen), and 75 μl PEI MAX. The mix was incubated at room temperature for 20 min before being added drop-wise to the culture plate while gently swirling it. The plates were incubated for 72-96 h. At the end of the incubation, the medium was removed and the cell monolayer was washed once in phosphate buffered saline, before scraping off the cells using a rubber policeman, and recovering them by centrifugation. Cell pellets were either snap frozen and stored at −80°C, or directly taken for purification of opsin.

For SiMPull and ensemble FRET experiments, opsin constructs with N-terminal HA-SNAP or C-terminal FLAG-SNAP were expressed in HEK293T cells or HEK293S GnTI^-^ cells and processed 24 h after transfection.

#### SiMPull experiments

SNAP-tagged opsins were labeled by incubating expressing cells with LD555 SNAP fluorophore. The cells were lysed with 1% (w/v) DDM for 1 h at 4°C. The lysate was clarified by centrifugation at 16,000 x g (4°C, 20 min) and diluted to 0.1% DDM for SiMPull. A glass flow chamber was prepared as previously described (20,21) using mPEG/biotin-PEG-passivated quartz slides and coverslips coated with neutravidin and subsequently decorated with biotinylated anti-HA antibodies. The flow chamber was placed on an inverted TIRF microscope (Olympus IX83 with cellTIRF system) and LD555-labeled HA-SNAP-opsin constructs were immobilized at a density that allowed clear resolution of individual spots (The fluorophore was excited using a DPSS 561 nm laser, imaged through a 100x objective (NA=1.49) and the data was collected at room temperature. Images were acquired with a scMOS detector camera (Hamamatsu ORCA-flash4.0 V3) at 20 Hz using Olympus cellSens software. Spots were analyzed using a previously described program (60) and manually characterized as bleaching in 1, 2, 3, or 4 steps (or deemed uncountable). Data in bar graphs represent averages of the step distributions for different movies.

#### Ensemble FRET experiments

Cells expressing SNAP-tagged opsin constructs were labeled with donor and acceptor fluorophores at a 1:2 molar ratio with SBG-(sulfonated, membrane-impermeant) or BG- (cell-permeable) derivatives of JF549 and JF646 dyes (28). Images were taken using a 60x objective on an inverted microscope using an scMOS detector camera (see SiMPull methods) in both donor and acceptor channels using donor excitation with low intensity 560 nm laser illumination before and after bleaching the acceptor with high-intensity 640 nm illumination. Regions of interest were manually drawn around the same clusters of cells and the percentage increase in donor fluorescence intensity following acceptor bleaching was calculated for at least 8 images per condition. Surface expression was calculated by measuring fluorescence intensity in the acceptor channel prior to bleaching using low intensity 640 nm illumination. For all comparisons the same laser intensities and exposure times (100 ms) were used.

#### Purification and quantification of opsin

Anti-FLAG-M2 agarose (~10 μl bed volume per 10-cm plate of cells) was pre-equilibrated by washing 3 times with 10 bed volumes of wash buffer (50 mM HEPES pH 7.4 100 mM NaCl, 1.0% (w/v) DDM). Cell pellets were incubated in 400 μl (per 10-cm plate of cells) of lysis buffer (10 mM HEPES pH 7.4, 100 mM NaCl, 10% (w/v) DDM, protease inhibitor cocktail) at 4°C for 40 min with end-over-end mixing. The lysate was clarified by ultracentrifugation at 150,000 x *g* for 20 min at 4°C, and the supernatant was transferred to the washed anti-FLAG-M2 agarose and incubated at 4°C for 2 h with end-over-end mixing. The resin was collected by centrifugation using a Pierce spin column and washed 3 times with 10 bed volumes of wash buffer, before incubating at 4°C for 1 h with wash buffer containing 0.15 mg/mL 3X-FLAG peptide (20 μl per 10-cm plate) to elute bound opsin. The eluate was collected by centrifugation, and the elution step was repeated. Purified opsin (typically a 4 μl aliquot) was analyzed by Coomassie-stained SDS-PAGE, in parallel with in-gel bovine serum albumin (BSA) standards (200-1000 ng). Band intensities were quantified by ImageJ and opsin was estimated in comparison with the linear calibration plot obtained with BSA standards. The yield of the purified opsin (similar for all constructs) was ~6 μg per 10-cm plate of cells, at a concentration of ~150 ng/μl. The purified protein was kept at 4°C for up to a week or snap-frozen in aliquots and stored at −80°C.

#### FRET measurements *in vitro*

WT and F220C opsins with a C-terminal FLAG-SNAP tag (16) were capture onto anti-FLAG resin and labeled with either SNAP-Surface 549 (donor (D)) or SNAP-Surface 649 (acceptor (A)). The resin was washed to remove unreacted dye, and the labeled proteins were eluted with FLAG peptide as described abovce. The labeled proteins (in 0.1% (w/v) DDM) were mixed in 1:1 ratio in a total volume of 400 μl. Samples are treated with 0 to 8 mg amounts of washed Bio-Beads SM2 for 1 hour at room temperature with end-over-end mixing. After the incubation period, the sample was removed and taken for fluorescence spectroscopy. Emission spectra (λ_ex_ =555 nm) were acquired for donor (D) alone, acceptor (A) alone, and D-A pairs of WT or F220C opsin. Buffer subtraction was done after first using the Rayleigh peak to scale the buffer spectrum to that of the sample (scale factor>1); the scaled buffer spectrum was used for subtraction. Buffer-subtracted spectra were further adjusted for protein loss due to BioBeads treatment by directly exciting the acceptor (λ_ex_ =650 nm), determining the area under the spectrum over the range 660-700 nm, calculating the ratio of the area under the spectrum for the D-A sample versus that of the corresponding A-only sample., and using this ratio as a correction factor. A FRET measure (FRET_*scaled*_) was obtained by calculating the ratio of the emission intensity at 670 nm (λ∈× =555 nm) for D-A mixtures relative to the A-only sample, followed by an offset correction corresponding to mock-treated samples.

#### Liposome preparation

A mixture of POPC, POPG (9:1) and NBD-PC (0.3 mol percentage) (total lipids 52.5 μmol) was dried in a round-bottomed flask using a rotary evaporator. The lipids were dissolved in diethyl ether and dried again in the rotary evaporator, followed by incubation in a vacuum dessicator overnight (or for at least 3 hr). The lipid film was rehydrated in 10 ml Buffer A (50 mM HEPES-NaOH, pH 7.4, 100 mM NaCl) by rotating (250 rpm) at 37°C for 20 min using the rotary evaporator (no vacuum) and then subjected to five freeze-thaw cycles (freezing in liquid nitrogen, thawing in a 30°C bath). The resulting suspension was extruded using a LIPEX extruder (Northern Lipids, Inc., Burnaby, BC, Canada) by 10 passes through 400-nm track-etch polycarbonate membranes followed by 4 passes through 200-nm track-etch polycarbonate membranes to generate unilamellar liposomes. The liposomes were kept at 4°C and used within 2 weeks. Phospholipids were quantified using a colorimetric assay as described previously (2,3), and the size distribution of the liposomes was determined by dynamic light scattering using a Litesizer 500 instrument (Anton Paar, USA).

#### Reconstitution of opsin - preparation of proteoliposomes

Unilamellar liposomes (800 μl, ~3.3 mM phospholipid) were destabilized in an 840-μl reaction containing 8.2 mM DDM in a 2-ml Eppendorf tube. After incubation for 3 h at room temperature (RT) with end-over-end mixing, DDM-solubilized opsin and Buffer A were added and the sample was mixed for an additional hour at room temperature. When titrating the amount of opsin to be reconstituted, the DDM levels of all samples was adjusted to that of the sample containing the maximum amount of opsin. To form sealed vesicles, DDM was removed by adding washed Bio-Beads SM2 as follows. First, 80 mg BioBeads were added, followed by end-over-end mixing for 1 h at RT. This was followed by the addition of 160 mg of beads and incubation for an additional 2 h at RT. The sample was then transferred to a glass screw-cap tube containing 160 mg of Bio-Beads SM2 and mixed end-over-end overnight at 4°C. We previously showed that the resulting proteoliposomes have undetectable levels of DDM, i.e. a DDM-to-phospholipid ratio of <1:100. The recovery of both opsin and phospholipids was ~65% as determined by quantitative western blotting and a colorimetric assay, respectively (3). Protein-free liposomes were reconstituted in parallel.

#### Scramblase activity assay

Liposomes or proteoliposomes (50 μl) were added to 1950 μL Buffer A in a plastic fluorimeter cuvette equipped with a stir bar. Time-based fluorescence was recorded (λ_ex_ = 470 nm; λ_em_ = 530 nm, excitation and emission slits 2.0 nm, data acquisition frequency 1 Hz, 25°C, stirring 900 rpm) in a temperature-controlled Horiba Fluoromax+ spectrofluorometer. After measurement for at least 50 s to ensure a stable signal, 40 mM dithionite (40 μl of a freshly prepared 1 M solution of sodium dithionite in unbuffered 0.5 M Tris) was added through the injection port in the lid of the fluorimeter chamber, and fluorescence was monitored for at least 500 s. Fluorescence traces were normalized to the starting value of fluorescence (F_0_) at the time of dithionite addition and analyzed by fitting to a double-exponential function. As described previously (3), the fast component of the fit with t_1/2_=16.8 ± 3.4 s (mean ± S.D. (n=27)), accounted for the majority of the fluorescence change in both liposomes and proteoliposomes. The t_1/2_ of the slow component was an order of magnitude greater; it’s origin is unknown (3) but may be due to a combination of dithionite entry and slow spontaneous scrambling that is present in all traces irrespective of the presence of reconstituted protein. The plateau value of fluorescence (F_min_) obtained from the fit was taken as the end-point of the assay and used to calculate the fraction of vesicles equipped with a functional scramblase (F_min_ = 0.55 for protein-free liposomes as previously reported (3)). Briefly, values of (0.55- F_min_) were calculated for all proteoliposome samples and normalized to the value obtained for samples reconstituted at a high protein-phospholipid ratio (PPR)(typically >0.48 mg opsin/mmol phospholipid) to obtain p(≥1) scramblase as described (3). PPR values were scaled to PPR* to include only those vesicles that could be reconstituted with protein. Thus, PPR*=PPR/x where x is twice the F_min_ value obtained for samples reconstituted at a high protein-phospholipid ratio (16). Plots of p(≥1) scramblase versus PPR* were fitted to a mono-exponential function (3,16), yielding a fit constant (a) that was used to compare the dimerization propensity of various opsin constructs.

## Data availability

All data generated or analyzed during this study are included in this published article and available from GK or AKM on reasonable request.

## Acknowledgements

We thank the Meyerson and Andersen laboratories (Weill Cornell Medical College) for a gift of HEK293S GnTI^-/-^ cells and use of a dynamic light scattering instrument, respectively, and Scott Blanchard (St. Jude Children’s Research Hospital, Memphis, TN) for the LD555 dye.

## Author Contributions

G.K., T.R.L., V.Y.A., J. Levitz and A.K.M. conceptualized the project. A.N.P. generated the opsin constructs, expressed, purified and reconstituted the proteins, and performed and analyzed scramblase assays. J. Lee performed ensemble FRET and SiMPull experiments. K.P. performed *in vitro* FRET experiments. G.K., A.M.P., Z.S. and M.A.C performed and analyzed MD simulations. A.K.M. oversaw the design and analysis of scramblase reconstitution and *in vitro* FRET experiments. G.K. oversaw the design and analysis of MD simulation experiments. J. Levitz oversaw the design and analysis of ensemble FRET and SiMPull experiments. G.K. and A.K.M. wrote the first draft of the manuscript. All authors edited the manuscript.

## SUPPORTING INFORMATION

**Figure S1:**
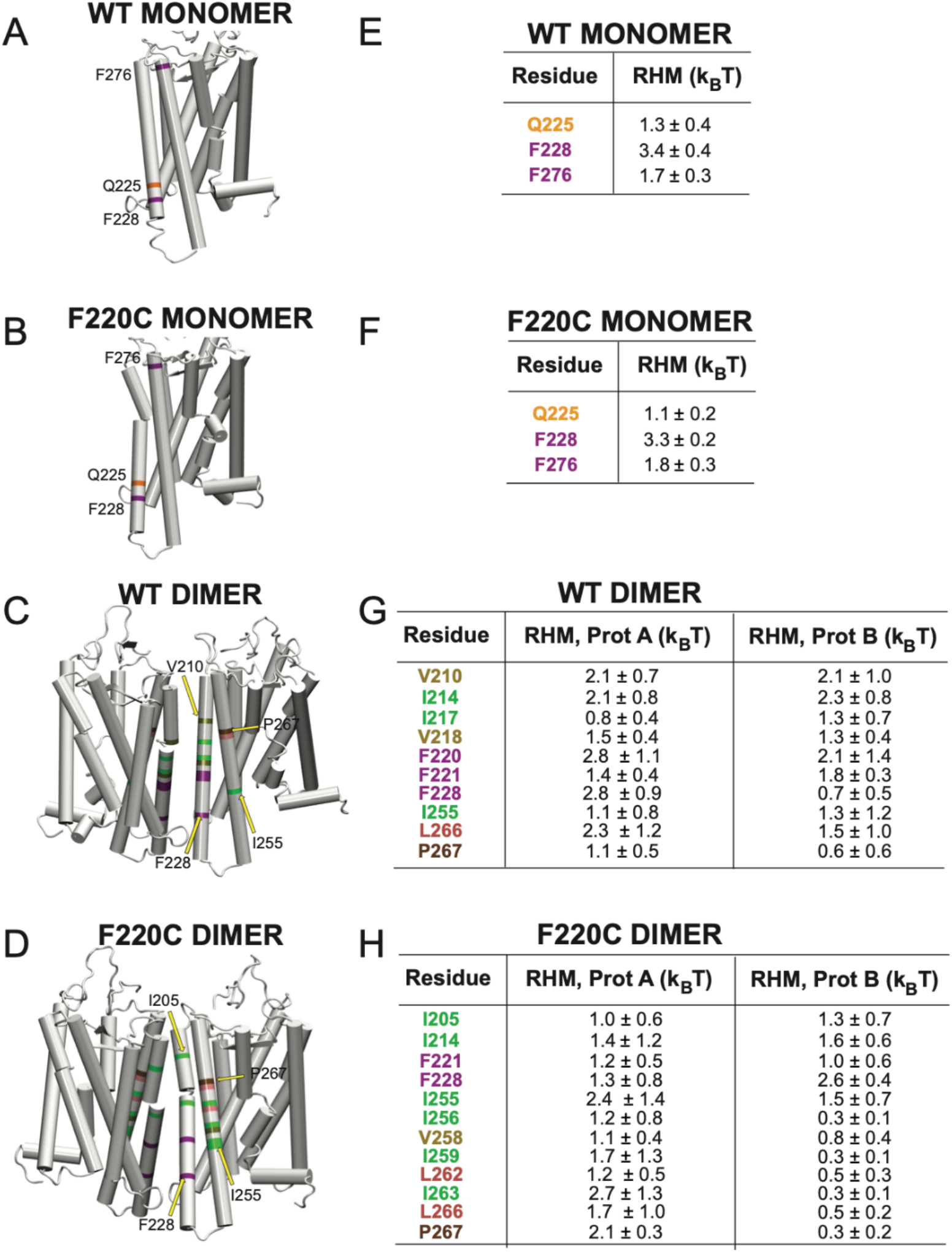
(**A-D**) Snapshots of WT and F220C monomer (A-B) and dimer (C-D) opsin constructs highlighting residues in TM5 and TM6 helices with high RHM penalty values. The residues are color coded based on their amino acid identity as following: Q – yellow, F – purple, I – green, L – pink, V – light brown, P – dark brown. Selected residues are labeled and marked with labels. (**E-H**) The RHM values for the residues highlighted in panels A-D. For the dimer systems, the RHM energies are separately shown for two the two subunits (Prot A, Prot B). Color code of the residues is the same as in panels A-D.

**Figure S2:**
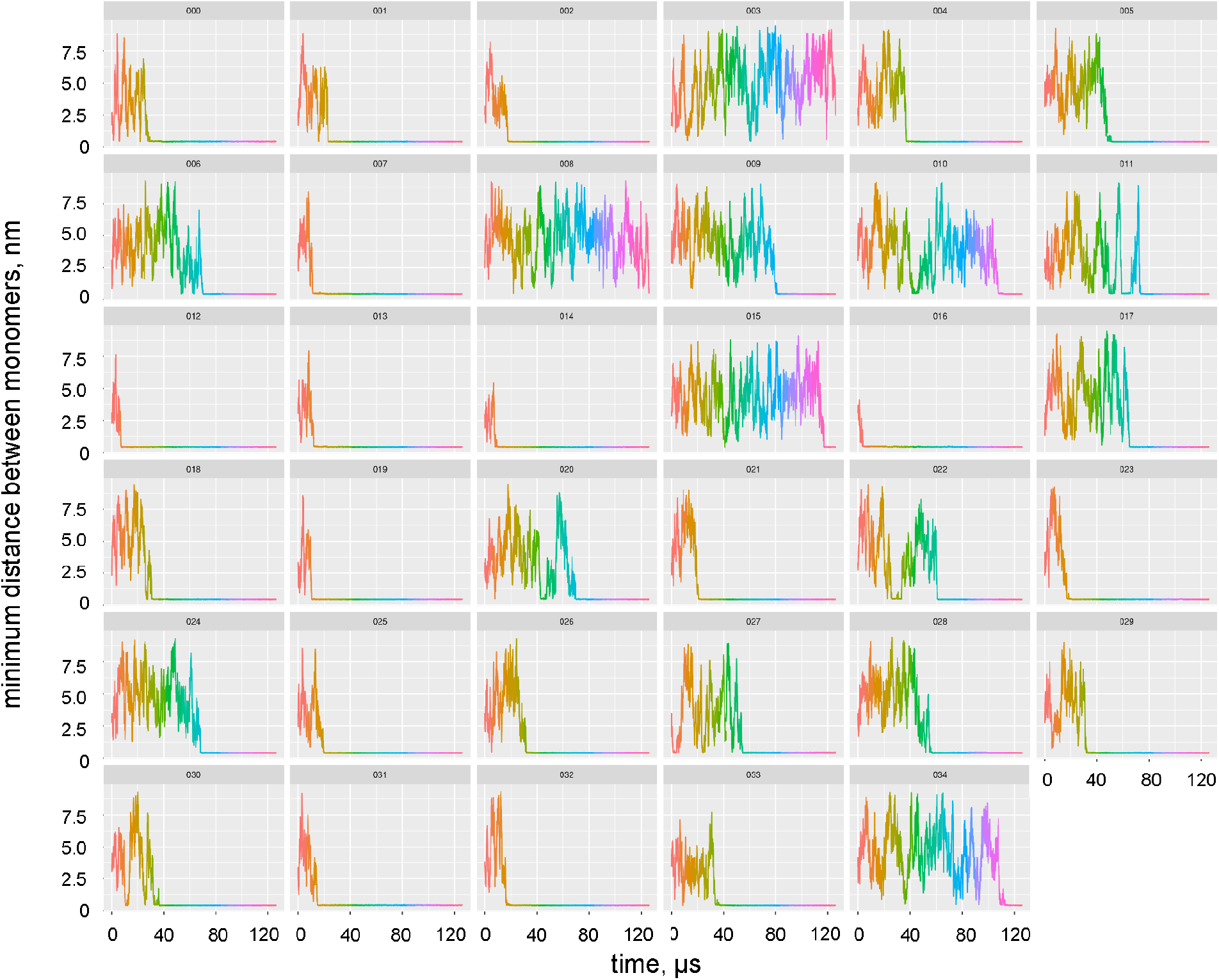
Time evolution of the minimum distance between two monomers of the WT opsin from the CG MD simulations. The data for each of the 35 replicates is shown in separate panels (labeled from 000-034). Small values of the distance reflect dimer formation. The rainbow color code in the panels, for clarity, represents simulation time.

**Figure S3:**
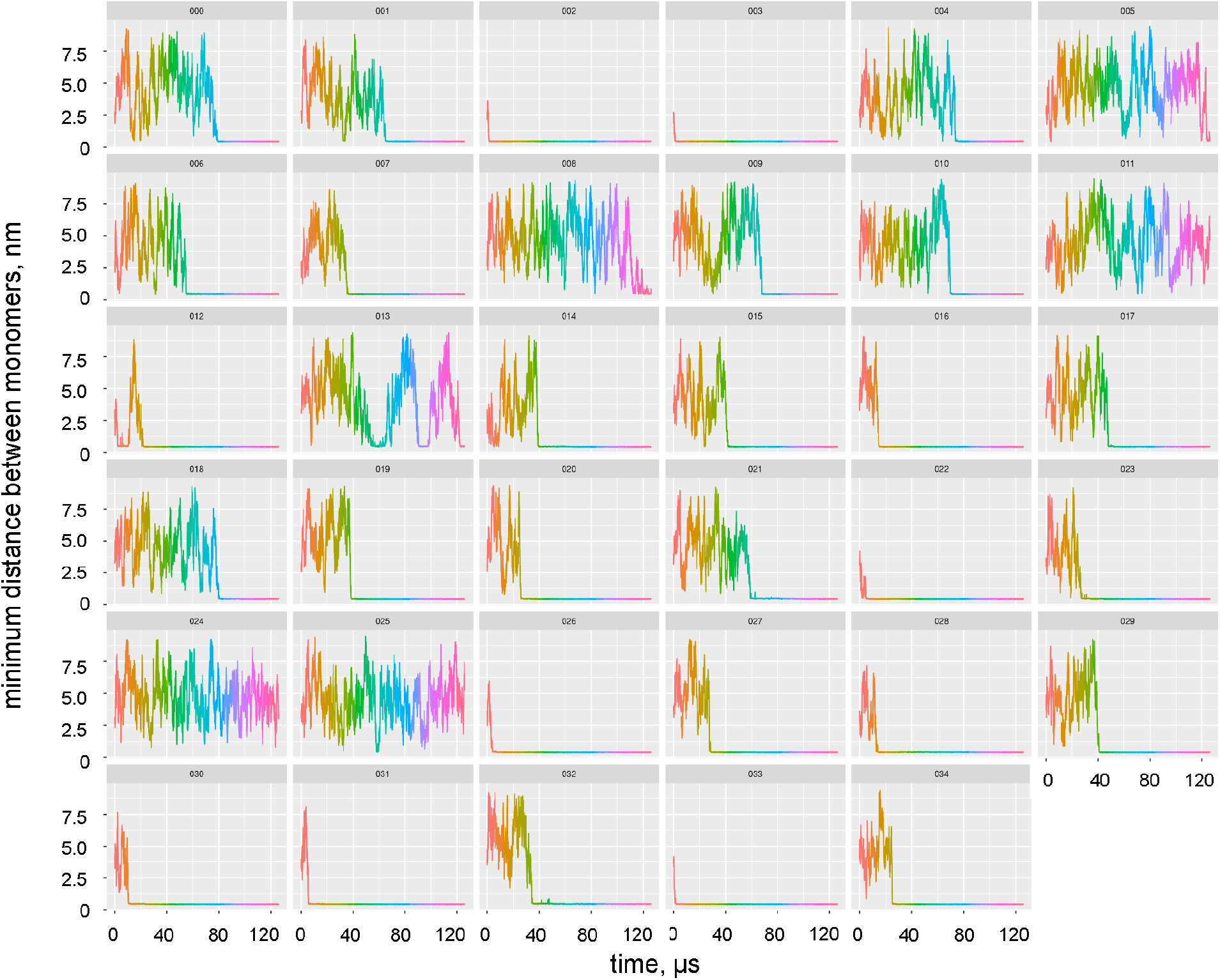
Time evolution of the minimum distance between two monomers of the F220C opsin from the CG MD simulations. The data for each of the 35 replicates is shown in separate panels (labeled from 000-034). Small values of the distance reflect dimer formation. The rainbow color code in the panels, for clarity, represents simulation time.

**Figure S4:**
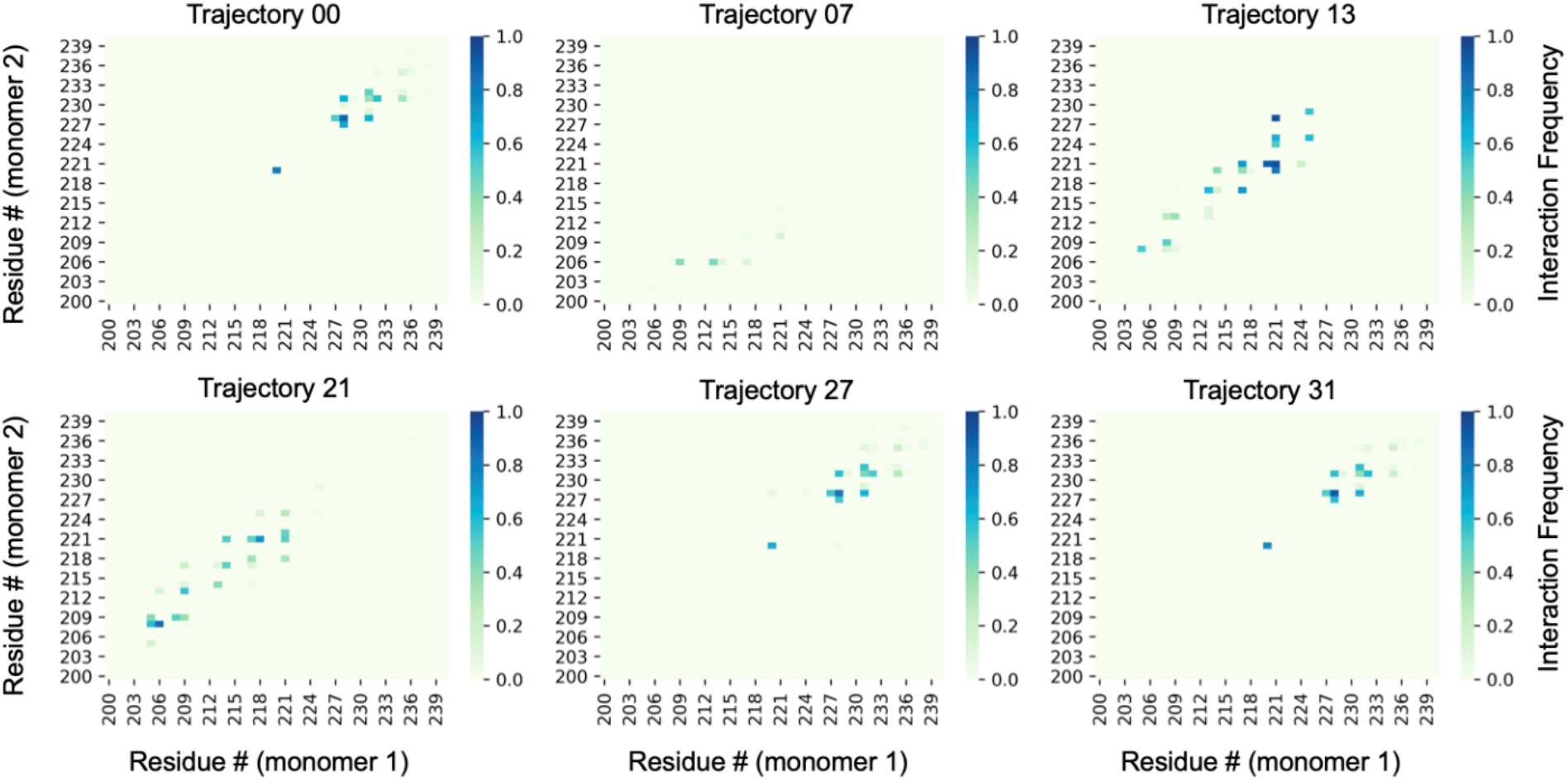
Pairwise residue contacts from the CG MD simulations of the WT opsin in which a dimerization interface was formed by interactions between TM5 segments on the opposite monomers of the dimer. The trajectory IDs are given on top of the panel. Color code represents normalized frequency of contacts between pairs of residues (see Methods for more details).

**Figure S5:**
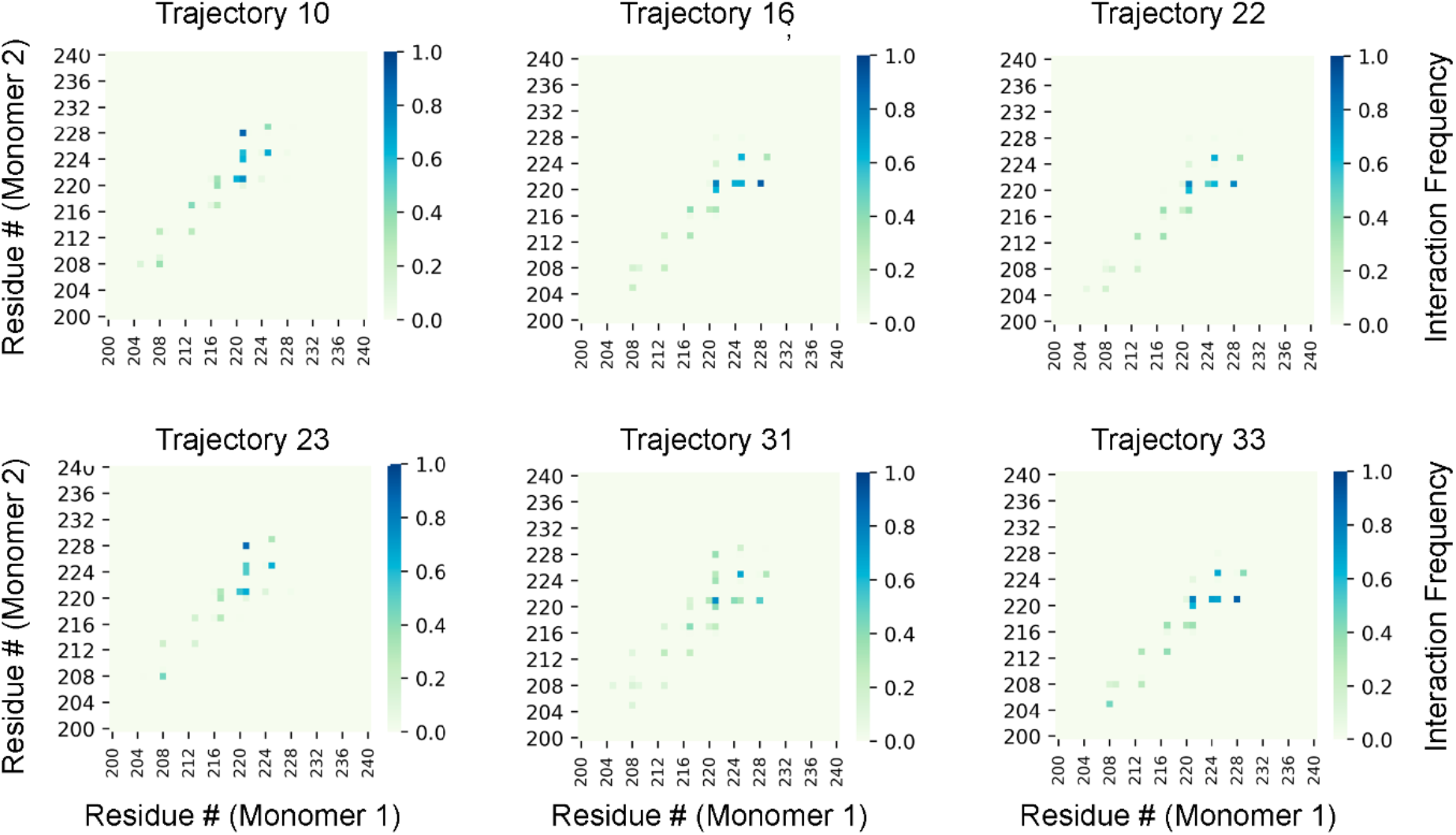
Pairwise residue contacts from the CG MD simulations of the F220C opsin in which a dimerization interface was formed by interactions between TM5 segments on the opposite monomers of the dimer. The trajectory ID-s are given on top of the panel. Color code represents normalized frequency of contacts between pairs of residues (see Methods for more details).

**Figure S6:**
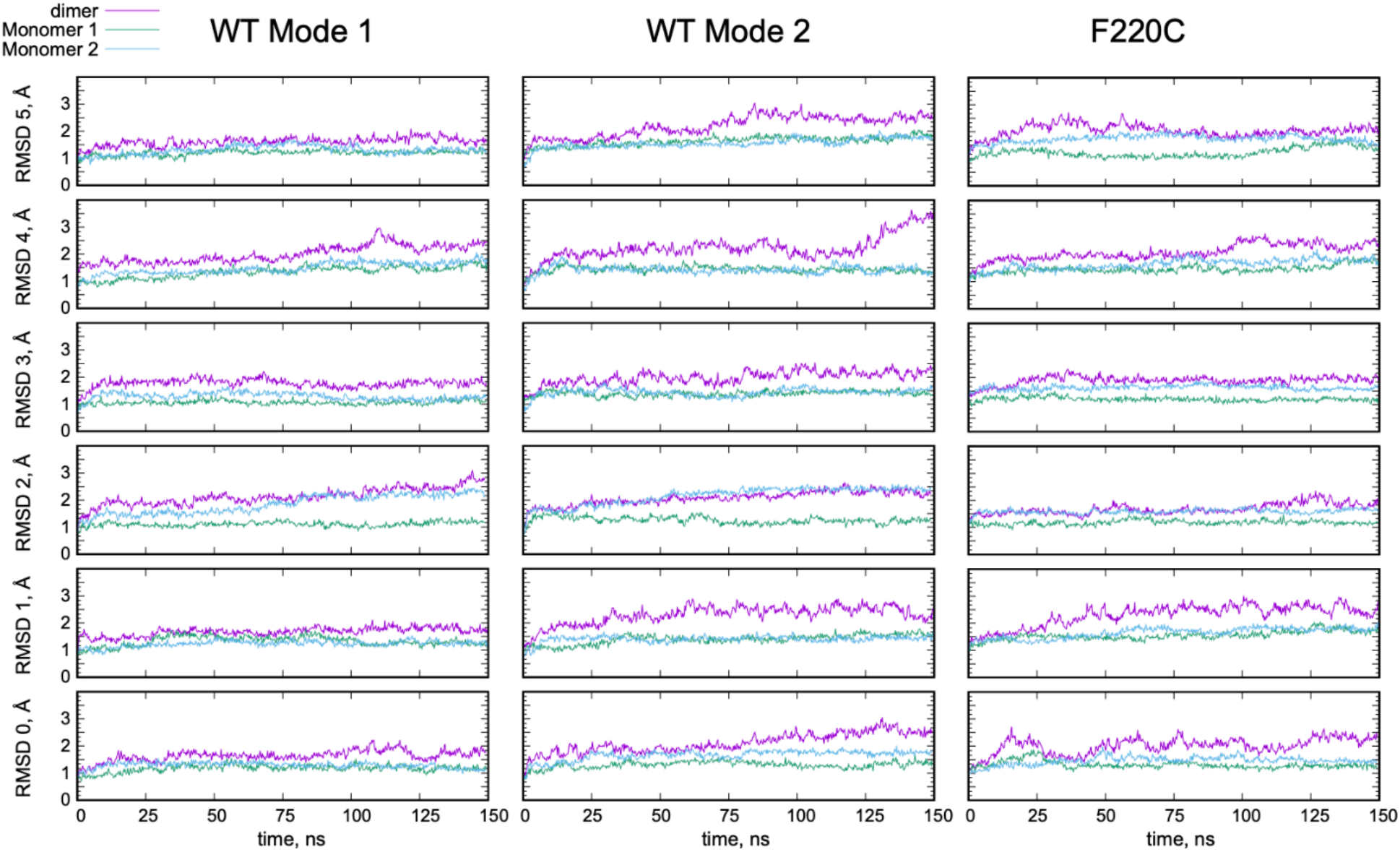
Time-evolution of root-mean-square deviation (RMSD) of the backbone atoms of the 7 transmembrane helical segments in the all-atom MD simulations of the Mode 1 and Mode 2 dimer models of the wild type opsin (WT Mode 1 and WT Mode 2), and of the F220C opsin dimer structure (F220C). The data for 6 independent replicates for each construct is given in separate rows. The RMSD was calculated for the entire dimer (purple traces), or for separate monomers (green and cyan traces). For each trajectory, the initial frame of the system was used as a reference structure for the RMSD calculations.

**Figure S7:**
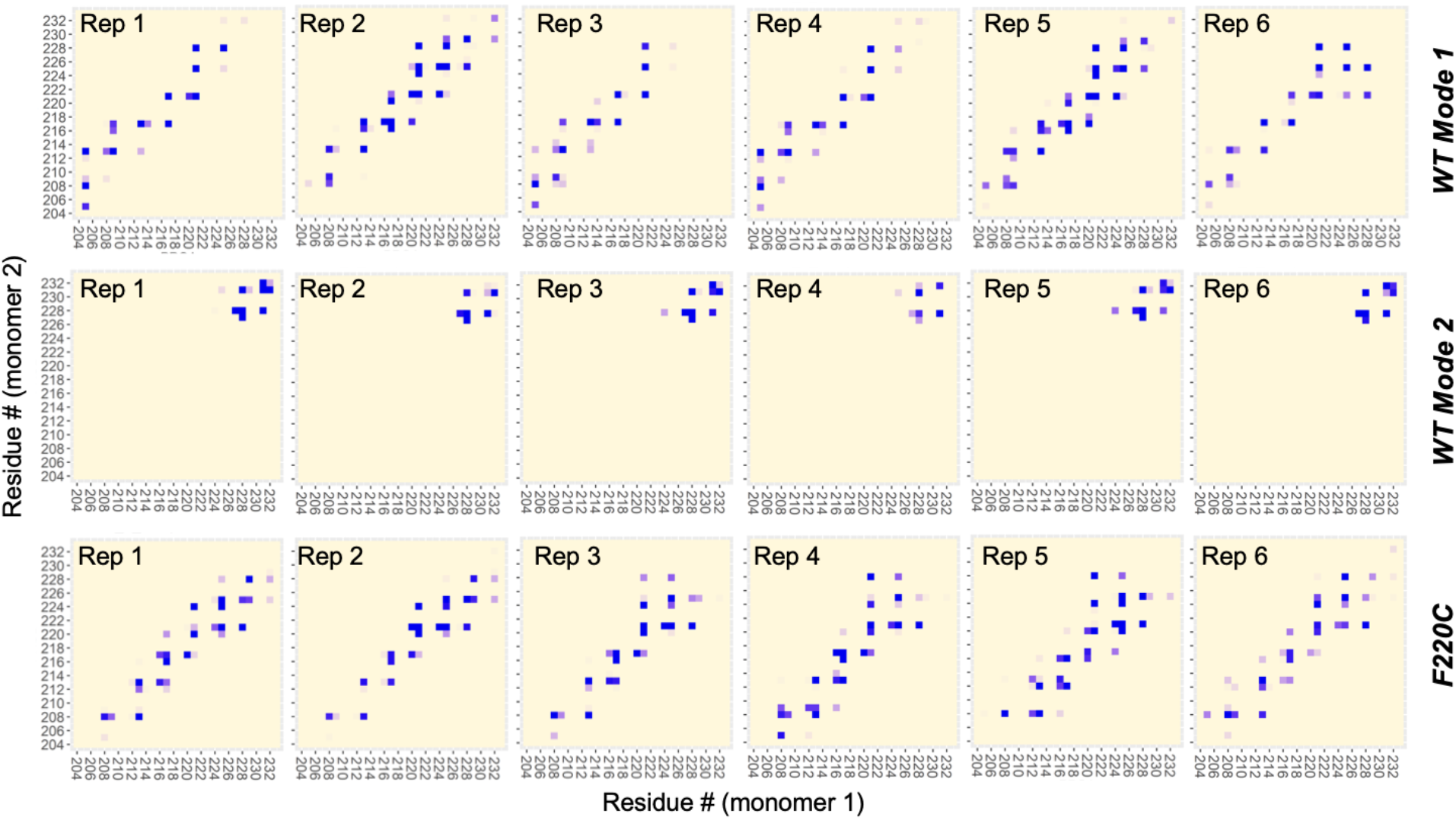
Pairwise residue contacts in the all-atom MD simulations of the two dimer models of the WT opsin (WT Mode 1 and WT Mode 2, top and middle rows) and of the F220C opsin dimer model (bottom row). The most frequent contacts are designated by dark blue shades, whereas the least frequent interactions are shown in light shades. The data for six independent replicates per construct is shown in separate panels, and the analysis was performed on the last 30ns of the all-atom trajectories.

**Figure S8:**
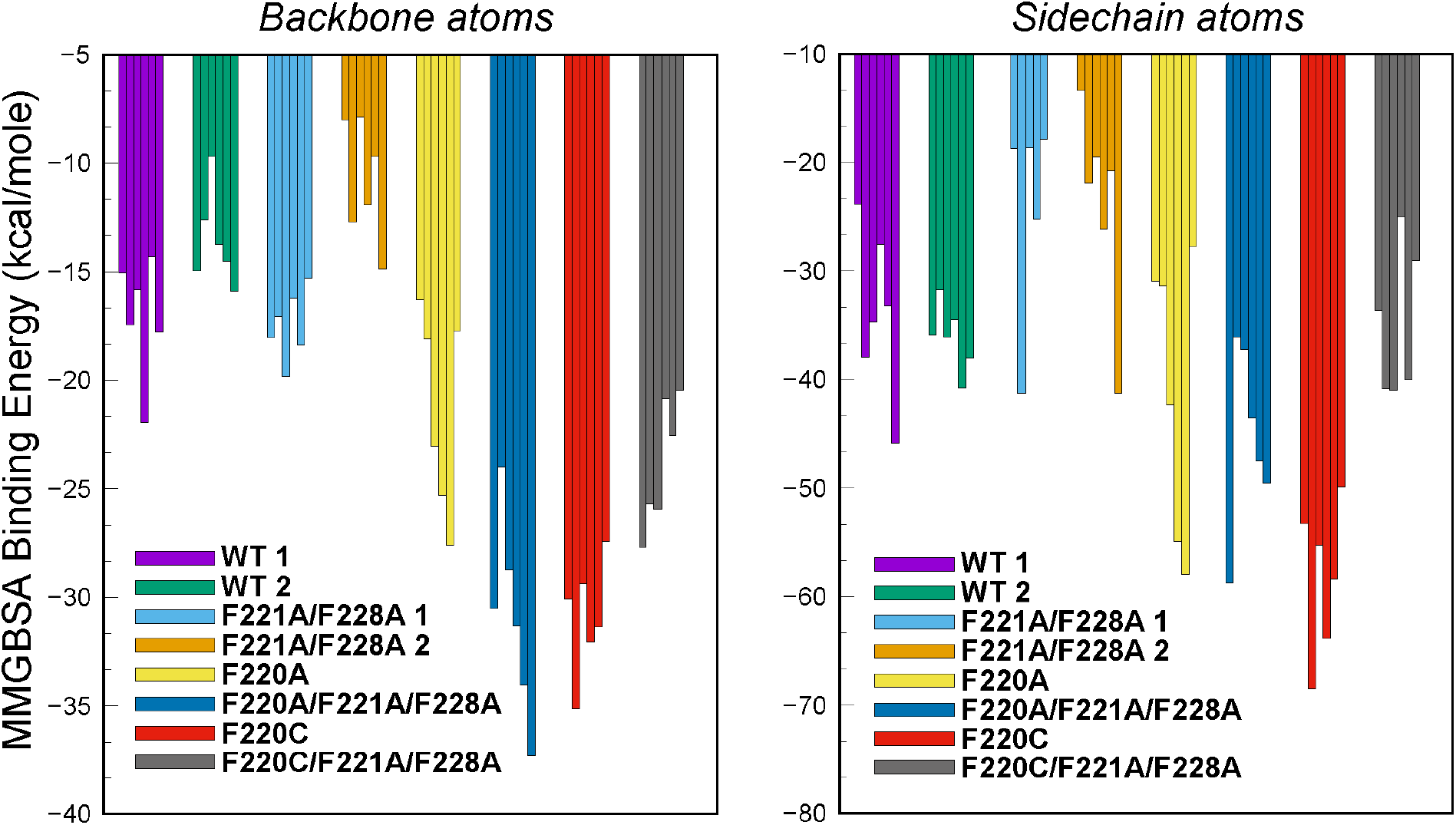
Contributions of the backbone (*left*) and sidechain (*right*) atom interactions to the overall MM-GBSA binding energy calculated from the all-atom MD simulations of the dimer models described in the text.

**Figure S9:**
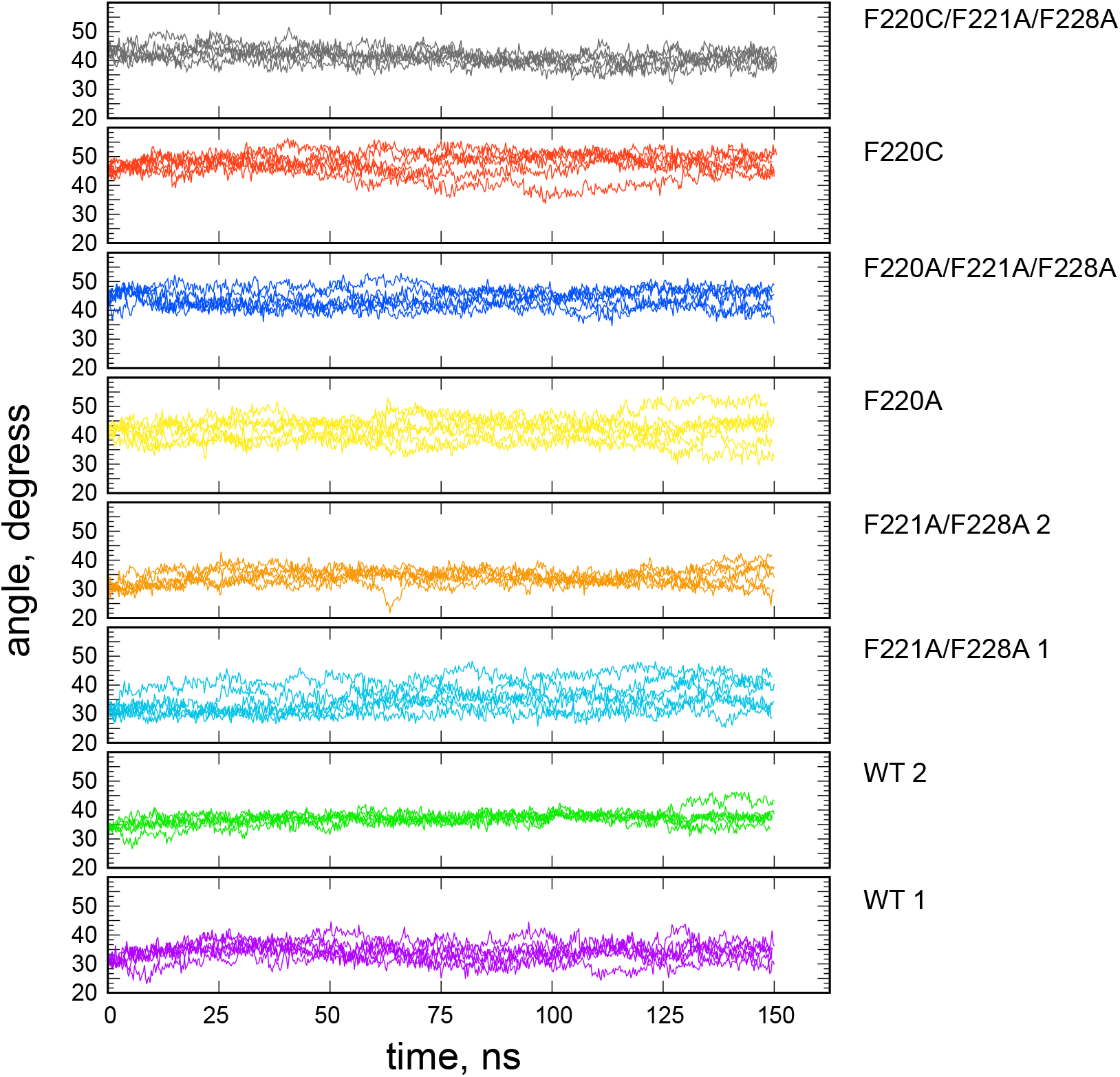
Time-evolution of the splay angle a in the all-atom MD simulations of the dimer models described in the text (see also Figure 7C in the main text). The color coding of the systems follows the same pattern as in Figures 6 and 7C in the main text. The results for each replicate are shown separately (i.e. 6 plots per constructs).

**Figure S10:**
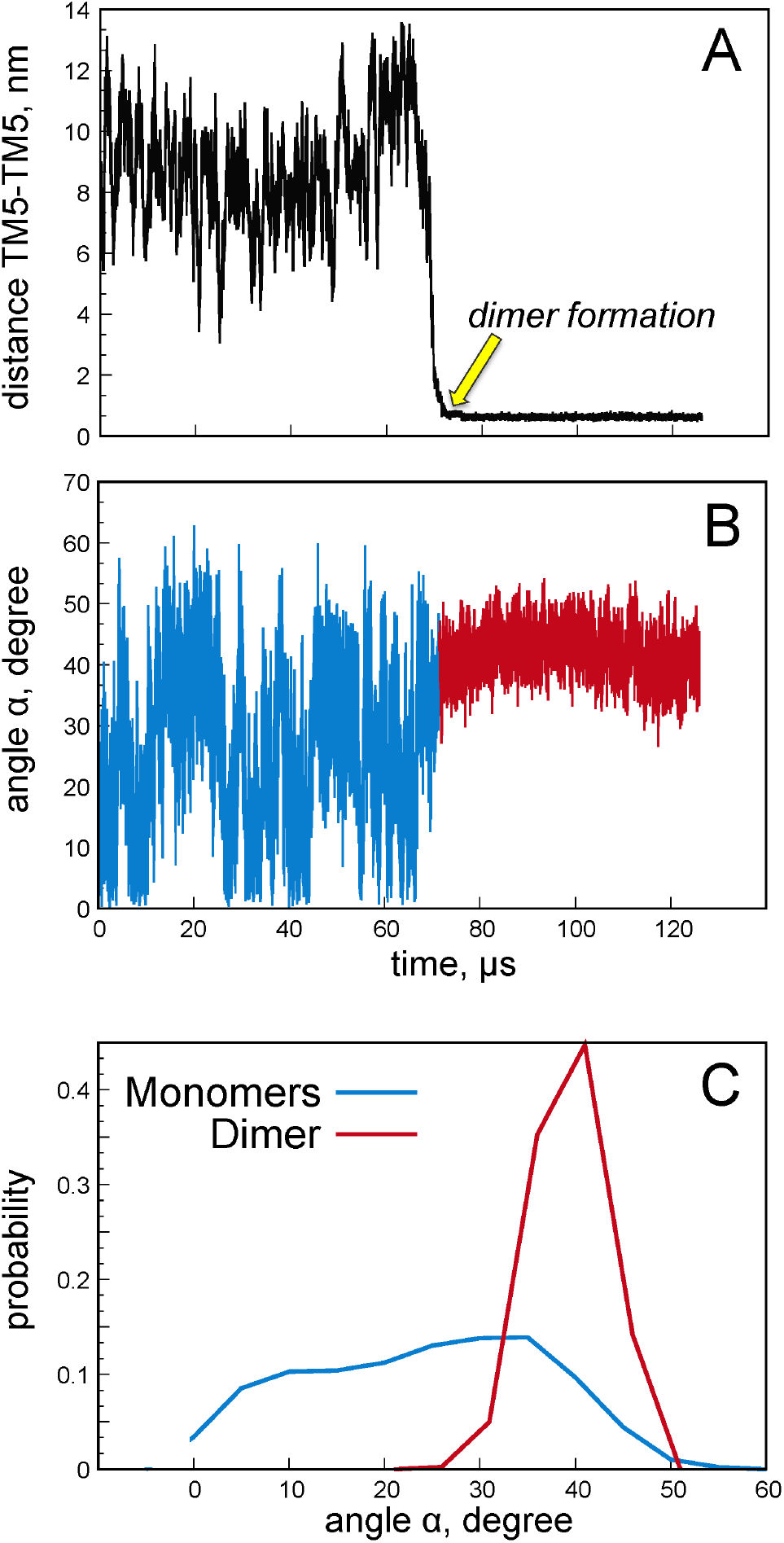
**(A-B)** Time evolution of the minimum distance between TM5 helices of the two monomers of opsin (A) and of the angle *a* between the two TM5 helices (B) in one of the CG MD simulations (Trajectory ID 10, see Figure S5). The time point of dimer formation is indicated with the yellow arrow. In panel B, the data for the trajectory parts before and after dimer formation are shown in blue and red colors, respectively. For definition of *a* angle see text. (**C**) Histograms of the *a* angle from panel B with the same color coding.

**Figure S11:**
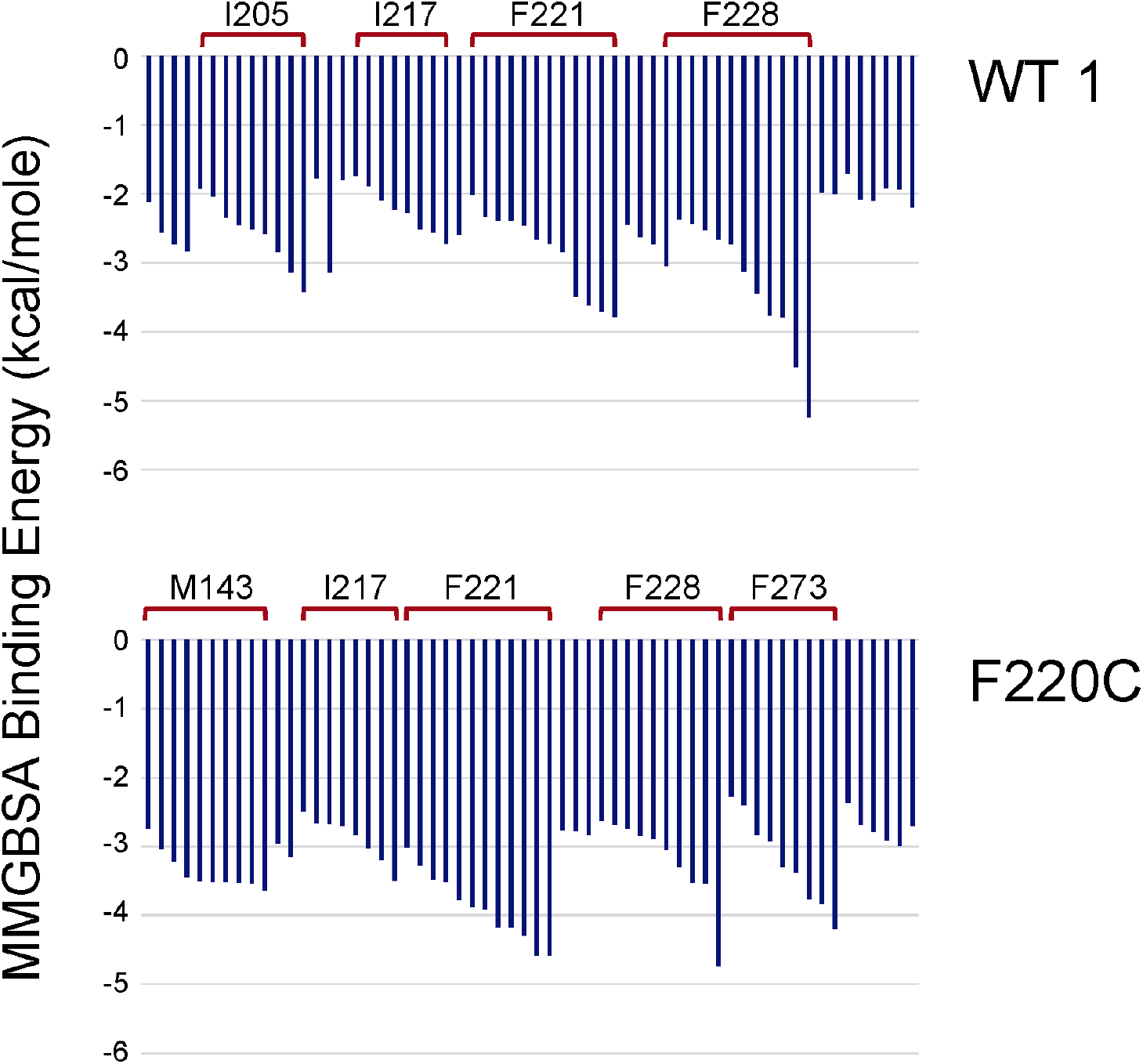
Per-residue decomposition of the MMGBSA binding energies for the WT Mode 1 and F220C dimer systems (upper and lower panels, respectively). Shown are contributions to the MMGBSA energy of the top 10 residue side-chains in all the trajectories for these two dimer constructs. The data is sorted according to residue ID from left to right. The data for the residues whose side-chains contributions were among top 10 in at least 4/6 trajectories are marked by red brackets and labeled.

**Figure S12:**
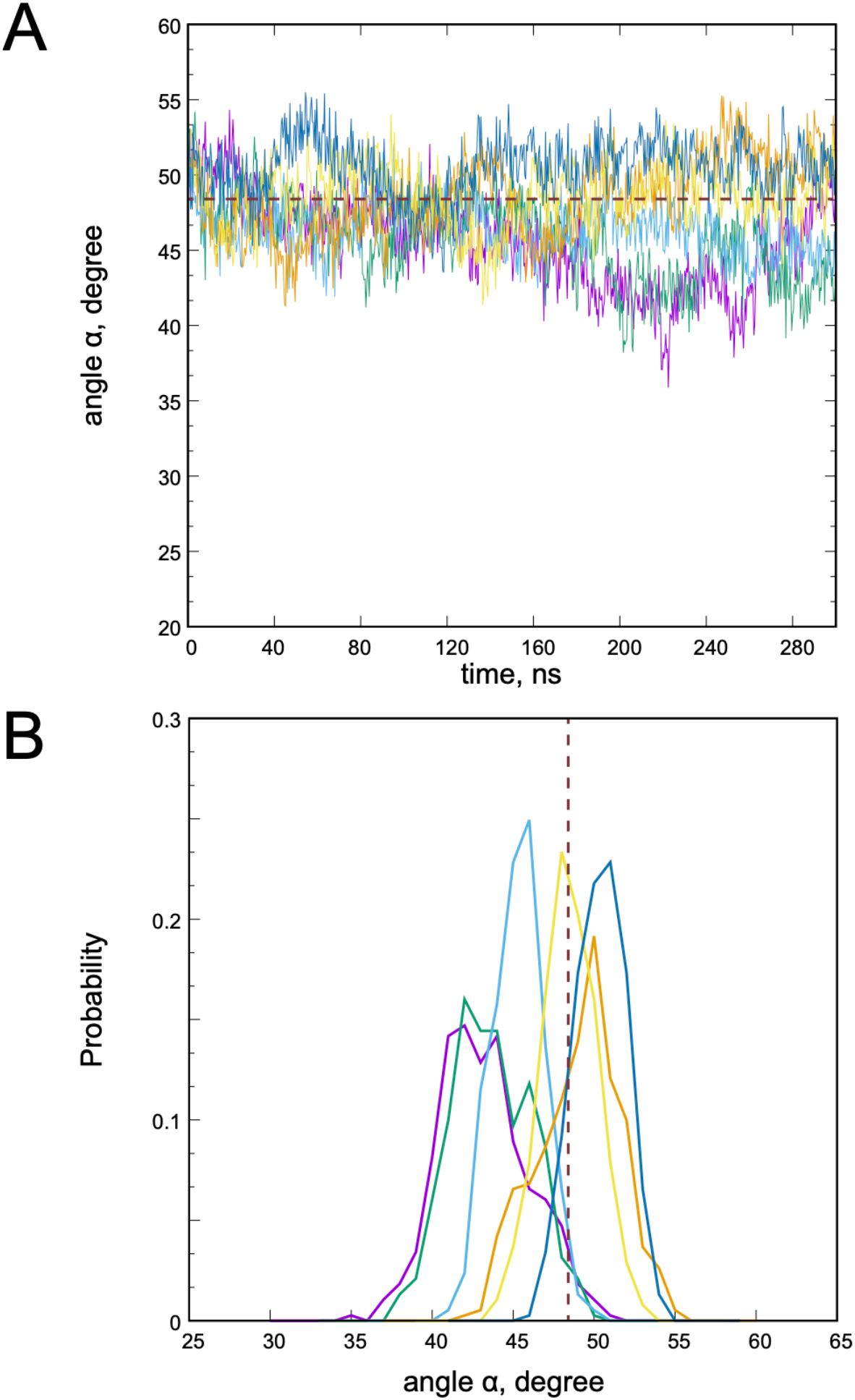
(**A**) Time-evolution of the α angle in the simulations of C220F opsin dimer. The data for 6 independent replicates are shown in different colors. The horizontal dotted line represents the value of the a angle (~48°) in the initial state of the system. (**B**) Histograms of the a angle from the panel A. The color scheme between the two panels is identical. The histograms were constructed from the analysis of the second halves of the MD trajectories described in panel A.

**Table S1:** Number of CG MD trajectories (out of 35 total replicates) of the WT opsin in which the dimerization was established via interactions between a specific pair of helical regions of the protein. ‘H8’ denotes helix 8 segment.

| Prot. A /B | TM1 | TM2 | TM3 | TM4 | TM5 | TM6 | TM7 | H8 |
| --- | --- | --- | --- | --- | --- | --- | --- | --- |
| TM1 | 6 | 6 | 0 | 0 | 3 | 2 | 1 | 1 |
| TM2 | 7 | 4 | 1 | 0 | 2 | 3 | 0 | 0 |
| TM3 | 7 | 4 | 0 | 0 | 1 | 0 | 0 | 1 |
| TM4 | 9 | 9 | 0 | 0 | 0 | 0 | 0 | 3 |
| TM5 | 8 | 4 | 2 | 1 | 6 | 5 | 0 | 0 |
| TM6 | 2 | 2 | 0 | 2 | 6 | 5 | 0 | 1 |
| TM7 | 2 | 3 | 1 | 1 | 0 | 0 | 0 | 0 |
| H8 | 1 | 0 | 0 | 0 | 3 | 2 | 0 | 4 |

**Table S2:** Number of CG MD trajectories (out of 35 total replicates) of the F220C opsin in which the dimerization was established via interactions between a specific pair of helical regions of the protein. ‘H8’ denotes helix 8 segment.

| Prot. A /B | TM1 | TM2 | TM3 | TM4 | TM5 | TM6 | TM7 | H8 |
| --- | --- | --- | --- | --- | --- | --- | --- | --- |
| TM1 | 8 | 8 | 3 | 2 | 5 | 3 | 0 | 2 |
| TM2 | 7 | 3 | 1 | 1 | 4 | 2 | 0 | 0 |
| TM3 | 4 | 1 | 0 | 0 | 6 | 0 | 0 | 2 |
| TM4 | 1 | 1 | 0 | 0 | 0 | 1 | 0 | 0 |
| TM5 | 4 | 3 | 6 | 0 | 7 | 5 | 0 | 3 |
| TM6 | 2 | 5 | 0 | 0 | 6 | 6 | 0 | 1 |
| TM7 | 1 | 1 | 0 | 0 | 0 | 0 | 1 | 0 |
| H8 | 3 | 0 | 1 | 2 | 3 | 0 | 0 | 6 |

**Table S3:** Listing of p-values between MMGBSA binding energies of various pairs of protein systems. The p-values were calculated from comparison of the mean and standard deviation values of the MMGBSA energies for each construct (see also Figure 6 in the main text). The abbreviations of the opsin constructs used in this table are as follows: WT 1 – WT Mode 1, WT 2-WT Mode 2, 2F/A – F221A/F228A, 3F/A – F220A/F221/F228A, C/2F – F220C/F221A/F228A. As highlighted in the red color, the dimerization energy for the F220C system is significantly different from the constructs that do not contain mutations at position 220, including the wild type dimers.

|  | WT 1 | WT 2 | 2F/A 1 | 2F/A 2 | F220A | 3F/A | F220C | C/2F |
| --- | --- | --- | --- | --- | --- | --- | --- | --- |
| WT 1 |  | 0.4 | 0.1 | 0.04 | 0.19 | 0.0014 | < 0.0001 | 0.0012 |
| WT 2 |  |  | 0.02 | 0.0045 | 0.33 | 0.0009 | < 0.0001 | 0.0005 |
| 2F/A 1 |  |  |  | 0.6 | 0.02 | 0.0002 | < 0.0001 | 0.0002 |
| 2F/A 2 |  |  |  |  | 0.01 | 0.0001 | < 0.0001 | 0.0001 |
| F220A |  |  |  |  |  | 0.0526 | 0.0015 | 0.0855 |
| 3F/A |  |  |  |  |  |  | 0.0335 | 0.63 |
| F220C |  |  |  |  |  |  |  | 0.0065 |
| C/2F |  |  |  |  |  |  |  |  |

